# Spatial organization of Dectin-1 and TLR2 during synergistic crosstalk revealed by super-resolution imaging

**DOI:** 10.1101/2022.04.25.489448

**Authors:** Miao Li, Christopher Vultorius, Manisha Bethi, Yan Yu

## Abstract

Innate immune cells recognize and elicit responses against pathogens by integrating signals from different types of cell-surface receptors. How the receptors interact in the membrane to enable their signaling crosstalk is poorly understood. Here, we reveal the nanoscale organization of TLR2 and Dectin-1, a receptor pair known to cooperate in regulating antifungal immunity, through their synergistic signaling crosstalk at macrophage cell membranes. Using super-resolution single-molecule localization microscopy, we show that discrete non-colocalized nanoclusters of Dectin-1 and TLR2 are partially overlapped during their synergistic crosstalk. Compared to when one type of receptor is activated alone, the simultaneous activation of Dectin-1 and TLR2 leads to a higher percentage of both receptors being activated by their specific ligands, and consequently an increased level of tyrosine phosphorylation. Our results depict, in nanoscale detail, how Dectin-1 and TLR2 achieve synergistic signaling through the spatial organization of their receptor nanoclusters.

## 1. Introduction

Innate immune cells, such as macrophage cells, use different types of receptors in combination to detect pathogens and elicit appropriate immune responses. In this process of receptor crosstalk, the immune response is orchestrated by two or more different types of receptors activated simultaneously. This combinatorial mechanism allows innate immune cells to achieve high detection specificity for a broad range of pathogens.^1-3^ While many pairs of immune receptors have been shown to function synergistically,^4-8^ the physical mechanisms by which different species of receptors interact during their crosstalk are poorly understood and, in some cases, controversial. The signaling crosstalk between Dectin-1 and Toll-like receptor (TLR) 2 is a hallmark example.

Both Dectin-1 and TLR2 are involved in regulating antifungal responses in innate immune cells. TLR2 recognizes lipopeptides on the surface of fungi and bacteria,^9, 10^ and Dectin-1 recognizes the β-1,3- and β-1,6-glucans in the cell wall structure.^11, 12^ Synergistic signaling crosstalk between the two receptors has been demonstrated by many studies. Dectin-1 signaling enhances TLR2-mediated proinflammatory immune responses in macrophages and dendritic cells, including the activation of nuclear factor kappa B (NF-κB)^13, 14^ and the production of cytokines.^15-17^ Meanwhile, TLR2 activation increases responses triggered by Dectin-1, such as the production of reactive oxygen species.^13, 18, 19^ But, while there is a consensus on the existence of Dectin-1 and TLR2 crosstalk, studies on just how Dectin-1 and TLR2 might interact to produce this synergy have yielded controversial results. In some studies, Dectin-1 and TLR2 were observed, in fluorescence microscopy observations, to co-exist on phagosome membranes, and co-immunoprecipitate in cell lysates. This led to the conclusion that the two receptors physically interact and colocalize spatially during their signaling crosstalk.^20-22^ However, some other studies reported that no direct physical interactions between Dectin-1 and TLR2 were detected, even though both receptors are concentrated on phagosomes that encapsulate zymosan particles ^13^. Recently, we reported, based on images acquired with total internal reflection fluorescence (TIRF) microscopy, that Dectin-1 and TLR2 form separate nanoclusters that do not colocalize but are within nanoscale proximity of one another.^23^ We showed that the strength of inflammatory responses from the TLR2 and Dectin-1 crosstalk requires this nanoscale proximity between both receptors.^23^ This study suggests that the spatial proximity of the receptors directly impacts their signaling crosstalk, but the means by which such crosstalk occurs, on the nanoscale, remain elusive.

Here we use two-color direct stochastic optical reconstruction microscopy (dSTORM) to resolve the nanoclustering of TLR2 and Dectin-1 during their synergistic signaling crosstalk in macrophage cell membranes. We found that TLR2 and Dectin-1 receptors each form nanoclusters that are only 100-200 nm in diameter. This is much smaller than what we reported before. This smaller size estimate is due to the improved spatial resolution of the current study. Importantly, we showed that Dectin-1 and TLR2 nanoclusters do not fully colocalize during synergistic signaling crosstalk, but are only partially overlapped. A larger percentage of Dectin-1 and TLR2 receptors are activated during synergistic crosstalk, than when either receptor is activated alone. This enhanced signaling competency of individual receptor nanoclusters leads to a higher level of tyrosine phosphorylation. Our results reveal the nanoscale organization of TLR2 and Dectin-1 during their crosstalk, providing detailed new clues about how this pair of receptors could interact to elicit enhanced inflammatory immune responses.

## 2. Materials and methods

### 2.1. Reagents

Reagents for ligand conjugation on glass coverslips include (3-aminopropyl) triethoxysilane (APTES, Sigma-Aldrich), poly-L-lysine (PLL) solution (0.1% w/v in water, Sigma-Aldrich), 1,1’-carbonyldiimidazole (CDI, Sigma-Aldrich), dimethyl sulfoxide (DMSO, Radnor), streptavidin (Invitrogen), Pam3CSK4 (Pam3, InvivoGen), biotin-labeled Pam3CSK4 (InvivoGen), beta-1,3-glucan from *Alcaligenes faecalis* (curdlan, InvivoGen), phosphate-buffered saline (PBS) (1×, pH 7.4, Sigma-Aldrich) and albumin from bovine serum (BSA, Sigma-Aldrich). Cytochalasin D (CytoD) was purchased from Enzo Life Science, Inc. Antibodies used for immunostaining include anti-CD282 (TLR2) (6C2) rat monoclonal antibody (eBioscience), anti-GFP goat polyclonal antibody, anti-MyD88 goat polyclonal antibody (R&D Systems, Inc.), anti-pSyk (Tyr525/526) (C87C1) rabbit monoclonal antibody (Cell Signaling Technology), AF488-labeled anti-phospho-tyrosine (pY) monoclonal antibody (pY20) monoclonal mouse antibody (Santa Cruz Biotechnology). Secondary antibodies, including sheep anti-goat antibody (Thermo Fisher Scientific), chicken anti-rat antibody (Thermo Fisher Scientific) and donkey anti-rabbit antibody (Thermo Fisher Scientific), were labeled with either Alexa Fluor 647 NHS Ester (Thermo Fisher Scientific) or Cy3B NHS Ester (Cytiva). The labeling efficiency was 2-3 dyes per antibody. Alexa Fluor 647 (AF647)-labeled phalloidin was purchased from Cell Signaling. Reagents used for dSTORM imaging include: 2-amino-2-(hydroxymethyl) propane-1,3-diol (Tris, Sigma-Aldrich), Buffer A (10 mM Tris, pH 8.0 and 50 mM NaCl), Buffer B (50 mM Tris-HCI, pH 8.0, 10 mM NaCl and 10% w/v glucose), 100×GLOX solution (3.4 mg/ml Catalase and 0.056 mg/ml glucose oxidase in Buffer A), and 1 M cysteamine (MEA) in water (pH 8.0).

### 2.2. Cells

RAW264.7 macrophage cells were purchased from American Type Culture Collection (ATCC) and cultured in Dulbecco’s modified Eagle medium (DMEM) supplemented with 10% fetal bovine serum (FBS), 100 U/mL penicillin, 100 mg/mL streptomycin, and 0.2 mM L-glutamine (Thermo Fisher Scientific). RAW264.7 macrophage cells stably expressing GFP-Dectin were originally made by Prof. David Underhill at Cedars-Sinai Medical Center. All cells were kept in an incubator at 37 °C with 5% CO_2_ and 95% relative humidity.

### 2.3. Functionalization of Pam3 and curdlan on glass coverslips

Round glass coverslips (35 mm in diameter) were first cleaned in piranha solution (3:1 H_2_SO_4_:H_2_O_2_, v:v) for 1 h at room temperature and dried with N_2_ stream. To conjugate curdlan on glass coverslips, the cleaned coverslips were aminated by incubating with 2% (v/v) APTES in anhydrous acetone at room temperature for 1 h, washed with pure ethanol, and then cured at 100 °C for 30 min. After cooling down at room temperature, the aminated coverslips were incubated with 0.5 M CDI in anhydrous DMSO at room temperature for 1 h to activate the primary amine groups. Afterward, coverslips were rinsed with DMSO and incubated with 1mg/mL curdlan in DMSO at room temperature for 2 h. Finally, coverslips were rinsed with DMSO and then PBS. The curdlan conjugated coverslips were stored in PBS at 4 °C. To prepare Pam3-coated glass coverslips, cleaned coverslips were incubated with 0.1% (w/v) PLL solution in water at room temperature for 1 h and rinsed with water. The PLL-coated coverslips were then incubated with 10 μg/mL biotinylated BSA in PBS at 4 °C overnight and rinsed with PBS. The biotin-BSA coated coverslips were then incubated with 5 μg/mL streptavidin in PBS for 1 h, rinsed with PBS, and incubated with 250 ng/mL biotinylated Pam3 (Pam3-biotin) at room temperature for 30 min. After the conjugation, the coverslips were rinsed with PBS and stored at 4 °C. To prepare glass coverslips functionalized with both Pam3 and curdlan, glass coverslips were functionalized first with curdlan and then with Pam3 following the procedures described above, except that streptavidin concentration was adjusted to 1.25 μg/mL. This adjustment was made to match the surface density of streptavidin on the bifunctional substrates to that on Pam3-only substrates.

In experiments to quantify the surface density of streptavidin on Pam3-only and Pam3-curdlan bifunctional substrates, AF647-streptavidin was mixed with unlabeled streptavidin at a molar ratio of 1:2000 during the conjugation. Using total internal reflection fluorescence (TIRF) microscopy, the total number of fluorescent streptavidin was counted using a single-particle localization algorithm^24^ and used to calculate the surface density of ligands.

### 2.4. Immunofluorescence staining

RAW264.7 macrophage cells stably expressing GFP-Dectin-1 were plated on glass coverslips at a density of ∼10^5^ cells/mL. After 10 min incubation (37 °C, 5% CO_2_), cells were washed with 1×PBS, fixed for 10 min with 4% (w/v) paraformaldehyde (PFA, Sigma) in 1×PBS, and then rinsed with 1×PBS three times for 5 min each. Cells were permeabilized with cold acetone (pre-chilled at -20 °C) for 10 min at -20 °C, rinsed in 1×PBS three times for 5 min each, and then passivated in blocking buffer (containing 1×PBS, 0.1% (v/v) Tween 20, 2% (w/v) bovine serum albumin (BSA), and 22.52 mg/mL glycine) for 1 h at room temperature. For immunostaining, cells were incubated with primary antibodies in the presence of 1% (w/v) BSA for 1h at room temperature, washed three times in washing buffer (containing 1× PBS, 0.1% Tween 20, and 1% BSA), and incubated with secondary antibodies in the presence of 1% (w/v) BSA for 1 h at room temperature. For immunostaining of GFP-Dectin-1 and TLR2, the primary antibodies consisted of anti-GFP goat IgG (5 μg/mL) and anti-TLR2 rat IgG (5 μg/mL), and secondary antibodies consisted of Cy3B-labeled sheep anti-goat antibody (2 μg/mL) and AF647-labeled chicken anti-rat antibody (2 μg/mL). For immunostaining of Dectin-1 and pSyk, primary antibodies consisted of anti-GFP goat IgG (5 μg/mL) and anti-pSyk rabbit IgG (5 μg/mL), and secondary antibodies consisted of Cy3B-labeled sheep anti-goat antibody (2 μg/mL) and AF647-labeled donkey anti-rabbit antibody (2 μg/mL). For immunostaining of TLR2 and MyD88, the primary antibodies consist of anti-TLR2 rat IgG (5 μg/mL) and anti-MyD88 goat IgG (5 μg/mL), and secondary antibodies consist of Cy3B-labeled chicken anti-rat antibody (2 μg/mL) and AF647-labeled sheep anti-goat antibody (2 μg/mL). For actin inhibition experiments, cells were seeded for 5 min and then incubated with 2.5 μg/ml cytoD for 5 min in an incubator (37 °C, 5% CO_2_). Cells were washed, fixed, permeabilized, and immunostained for GFP-Dectin-1 and TLR2 following the protocol described above.

### 2.5. Epi-fluorescence and total internal reflection fluorescence (TIRF) microscopy

All epi-fluorescence and TIRF imaging were performed using a Nikon Eclipse Ti2 inverted microscope equipped with a 1.49 NA ×100 oil objective and an Andor iXon3 electron-multiplying charge-coupled device (EMCCD) camera.

### 2.6. Direct Stochastic Optical Reconstruction Microscopy (dSTORM) and data analysis

1. Image acquisition. dSTORM imaging was performed in TIRF mode using a Nikon Ti2 Eclipse microscope equipped with a perfect focus system (PFS), a TIRF ×100 oil-immersion objective (NA 1.49) and an Andor iXon3 EMCCD camera. Dual-color dSTORM images were acquired sequentially under the excitation of 637 nm and 561 nm diode lasers at a power density of 10 and 8 kW/cm^2^, respectively. Images were acquired with an exposure time of 10 ms for 10,000 consecutive frames per fluorescence channel. A 405 nm diode laser at a power density of 0.5 kW/cm^2^ was used to stochastically activate fluorophores. Fixed and immunostained cell samples were kept in freshly made dSTORM imaging buffer (100 mM MEA and 1×GLOX solution in Buffer B) and imaged at room temperature.
2. Image post-processing. dSTORM images were reconstructed using ImageJ plugin ThunderSTORM^25^. A multiple-emitter fitting analysis (MFA)^26^ was used to resolve single molecules in high spatial densities. Several filters were applied sequentially to remove noise in images.
  i. To remove molecules with low photon detections, an intensity filter with a threshold of *N* >*αb* was applied, where *N* is the number of photons detected from a given molecule; *b* is the standard deviation of the signal in the fitting region, and α, a sensitivity factor (typically ranging from 20 to 40) was set as 30 in this study^27^.
  ii. To remove the noise generated from non-clustered molecules, a local density filter was applied. At each localization, the number of its neighboring molecules at a given radius was counted, and if below a given threshold, that localization was discarded. In this study, a minimum number of 10 neighbors in the radius of 60 nm was chosen as the threshold to effectively remove the non-clustered molecules.
  iii. To eliminate multiple blinking from a single molecule in sequential frames, localizations identified within a single pixel of the CCD camera (105 nm here) in five consecutive frames were considered as the same localization and merged into a single localization by averaging the coordinates from all localizations to be merged. The photon number and the background signal level of the new merged localization were calculated as the sum values of those parameters from the pre-merged localizations. The inaccuracy of the new merged localization was calculated as:

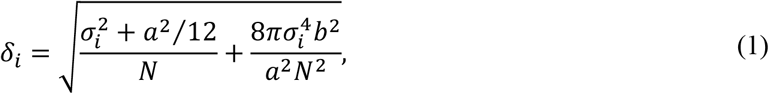

where *N* is the number of photons collected from a given molecule, *σ*_*i*_ is the fitted standard deviation of point-spread functions (PSF) of the detected molecules in either the x or y direction, *a* is the pixel size of the CCD camera and *b* is the standard deviation of background.
  iv. To remove repeated localizations of a single molecule in one frame, such as when multiple dyes on a single protein blink simultaneously and are detected, localizations that were within the scale of the localization inaccuracy in one frame were grouped, and in each group, only the molecule with smallest localization inaccuracy was kept. Two-color dSTORM images were processed for each channel separately following the same procedure, and the lateral drift between two fluorescence channels was corrected using TetraSpeck fluorescent particles (100 nm in diameter) that were added in the cell samples as fiducial markers during dSTORM imaging. Because two-color dSTORM imaging was performed sequentially in 637 nm excitation channel first and then in 561 nm channel, the first frame in the 637 nm channel was used as reference to correct the lateral drift of both channels. Briefly, the x-y coordinates of individual fiducial marker (*x*_*m,N*_ and *y*_*m,N*_) in each frame in 637 nm channel were averaged as:

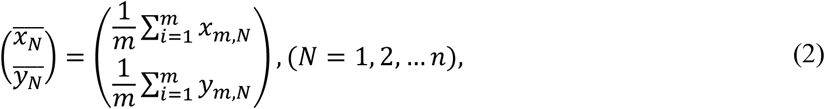

where *m* is the number of fiducial markers in each frame, *N* is the total frame number. The averaged coordinates of fiducial markers 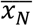 and 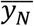 in each frame n were then compared to that in frame 1 to obtain the average displacement of each frame relative to frame 1:

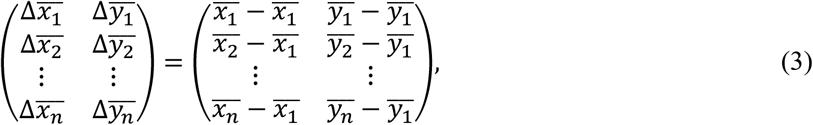

The relative averaged displacement 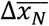 and 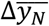 frame n were then subtracted from every single-molecule localization *X*_*n*_ and *Y*_*n*_ in the corresponding dSTORM image of the cell sample, generating the corrected x-y localization 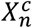 and 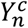 in each frame:

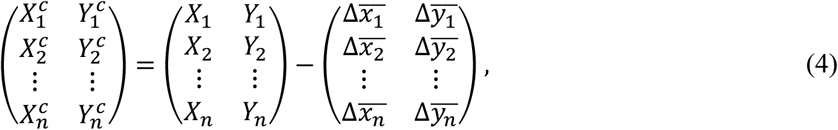

where *X*_*n*_ and *Y*_*n*_ represent all the original x and y coordinates of the cell sample in each frame n. 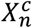 and 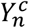 represent all the corrected x-y coordinates from each frame n. The same procedure was applied to correct lateral drift in images from the 561 nm excitation channel using frame 1 in 637 nm channel as a reference. Τhe final dSTORM images were visualized using Gaussian rendering or single-molecule localization scatter plot. Gaussian rendering was achieved using ImageJ plugin ThunderSTORM with a standard deviation equal to the localization inaccuracy. Single-molecule localization scatter plot was generated using R package “ggplot2”^28, 29^.
3. Receptor cluster analysis. After post-processing of dSTORM images, single-molecule localizations of TLR2 and Dectin-1 were grouped into nanoclusters using Topological Mode Analysis Tool (ToMATo),^30^ a persistence-based clustering segmentation algorithm.^31^ Briefly, the algorithm first estimates the detection density at each single-molecule localization by counting the number of its neighboring detections within a circle with a fixed search radius *r*. The candidate clusters were formed using a mode-seeking approach. This algorithm links each single-molecule localization to its neighboring localizations having the highest detection density within a circle of a search radius *r*. All the linked localizations then formed a candidate cluster. Each candidate cluster has a maximum point in density and a saddle point, which are known as the birth density and death density, respectively. The difference between the birth and death density is defined as the persistence of the candidate cluster. A persistence threshold was chosen using a persistence diagram. This diagram consists of the scatter plot of the birth density and death density of each candidate cluster in a Cartesian coordinate system. The intersection point of the x axis (birth density) with line y=x-*τ* was recorded, where *τ*, the x axis value at that intersection point is the persistence threshold. Candidate clusters with persistence below the persistence threshold were merged to neighboring candidates that have persistence greater than, or equal to, the threshold. In this study, a search radius *r*=25 and a persistence threshold *τ*= 6 were chosen based on a previously reported simulation for the performance of ToMATo across all tested parameters.^30^ These optimized values resulted in a high-performance cluster detection (>90% accuracy).^30^ Following the identification of clusters, the ToMATo clustering analysis was performed using the R package “RSMLM”^30^. The convex hull of all single-molecule localizations within a cluster was obtained as the cluster area (Fig. S4C). The diameter of a cluster was calculated as the average of the major and minor axes of the bounding ellipse of each cluster (Fig. S4D). To estimate the bounding ellipse, a customized written R code, based on Matlab code by N. Moshtagh,^32^ was used to calculate an ellipse of a minimum area enclosing all of singe-molecule localizations within a cluster. To optimize the parameters of identifying the receptor nanoclusters, dSTORM images of secondary antibodies that were non-specifically adsorbed in cells were used as controls. In those control images, the number of detected nanoclusters per unit area and the number of single-molecule localizations per nanocluster were quantified (Fig. S5). In this study, we found a mode value of 15 localizations per receptor nanocluster (both TLR2 and Dectin-1) and a mode value of 8 localizations per antibody nanocluster (both 637 nm and 561 nm excitation channel). Therefore, we used “localizations ≥ 15” as a minimum number of localizations per receptor nanocluster, which could minimize the possibility of recognizing non-specifically adsorbed antibodies as receptor nanoclusters.
4. Analysis of cluster overlap. The degree of overlapping between two types of protein nanoclusters was estimated using a coordinate-base1d colocalization analysis^33^ available in the ImageJ plugin ThunderSTORM. Briefly, for each single-molecule localization (*A*_*i*_) of protein A, the density of localized single molecule localizations of protein A and that of protein B around *A*_*i*_ within a given circle of radius *r* were calculated as:

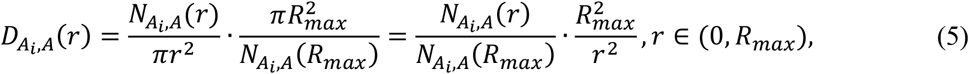

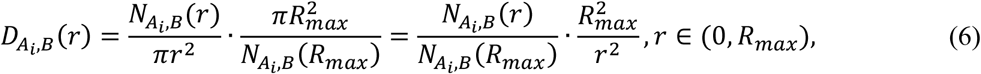

where 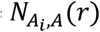 is the number of localizations of protein A within a circle of radius *r* around *A*_*i*_ and 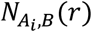 is the number of localizations of protein B within a circle of radius *r* around *A*_*i*_, *R*_*max*_ is the maximum radius set by the user. The two density gradients 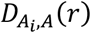 and 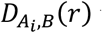 were compared by calculating their Spearman correlation coefficient 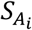. Then, the degree of colocalization (DoC) score, 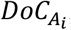, was calculated as:

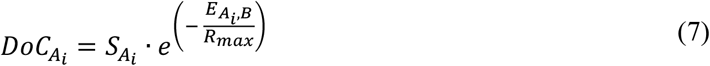

where 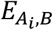 is distance from *A*_*i*_ to its nearest neighbor protein B. As such, each single-molecule localization in the two-color dSTORM images was assigned a DoC score ranging from −1 (anti-colocalized), through 0 (no colocalization), to +1 (perfectly colocalized) (Fig. S9). In this study, we performed the CBC analysis for the single-molecule localizations between TLR2 and Dectin-1, MyD88 and pSyk, TLR2 and MyD88, as well as Dectin-1 and pSyk at increasing radii using the following parameters: search radius of 10 nm, radius step of 10 nm, and step count of 25. Single-molecule localizations with DoC scores ≥ 0.4 were recognized as overlapping. The total number of overlapping localizations divided by the total number of localizations in each species of proteins was calculated as the degree of overlapping.

The accuracy of this overlapping analysis and the determination of using a DoC threshold value of ≥0.4 was validated by simulations (Fig. S12) following the previously reported procedure.^34, 35^ First step in the simulation was to create two sets of identical nanoclusters with varying degree of overlapping. To create this, a cluster of localizations of 35 nm in diameter was selected in dSTORM images and the single-molecule localizations inside this nanocluster were duplicated and then translocated 30 nm away from its original position, resulting in an overlapping region between two identical nanoclusters. DoC scores were calculated using parameters: search radius of 10 nm, radius step of 10 nm, and step count of 25. A localization inaccuracy of dSTORM imaging (approximately 20 nm; Fig. S3) was applied to indicate the errors of the analysis (Fig. S12). To test how the size of nanoclusters affects the accuracy of the overlapping analysis, a larger receptor nanocluster of 182 nm in diameter and molecule density of 0.009 (# localization per unit area of a cluster) was duplicated and laterally translocated to vary the degree of overlapping, from complete overlapped through partially overlapped to separated (Fig. S14). DoC scores were calculated and examined using cluster maps and histograms (Fig. S14). To test how the molecular density of nanoclusters affects the analysis accuracy, a denser receptor nanocluster (diameter 218 nm; molecule density 0.085) was duplicated and laterally translocated as described above (Fig. S15). DoC scores were calculated and examined using cluster maps and histograms (Fig. S15).

### 2.7. Analysis of pY intensity from fluorescently immunostained samples

To measure the fluorescence intensity of individual pY puncta, the centroid of individual pY puncta and the full-width-at-half-maximum (FWHM) of the point-spread function of each puncta fluorescence intensity profile were identified using a single-particle localization algorithm.^24^ From the single-particle tracking algorithm, the number of pixels (size) and the integrated intensity of each punctum were obtained. The fluorescence intensity of a single punctum was calculated as the intensity per pixel, which is the integrated intensity divided by the number of pixels of the punctum. To measure the pY intensity from each cell, a mask that differentiates individual cells based on pY intensity threshold was created. The pY intensity per unit area within that mask was calculated as the integrated pY intensity divided by the total number of pixels within that mask.

### 2.8. Statistical analysis

Statistic figures were plotted using both Prism 8 Software (GraphPad, USA) and R package “ggplot2”.^28, 29^ Data were shown as average mean ± standard error of the mean (SEM). Mann-Whitney U test was performed for two-group comparisons. Kruskal-Wallis test with Dunn’s test as a post-hoc test was performed for multiple-group comparisons. Statistical significance is indicated as follows: not significant (ns) p > 0.05; significant: * p <= 0.05; ** p <= 0.01; *** p <= 0.001; **** p <= 0.0001

## 3. Results and discussion

### 3.1. Nanoclustering of activated TLR2 and Dectin-1 in macrophage cell membrane

RAW264.7 macrophage cells stably expressing Dectin-1 green fluorescent protein (GFP) were used in this study, as wild-type RAW264.7 cells express only a negligible level of Dectin-1.^20, 36^ The immune responses of these cells were activated by ligands coated on glass coverslips. The Lipopeptide Pam3CSK4 (Pam3) was used as a ligand for TLR2, and the β-1,3-glucan polymer curdlan for Dectin-1. We prepared three different kinds of ligand-coated coverslips. The first kind had a substrate containing only curdlan, intended to activate only Dectin-1. To prepare such coverslips, we covalently conjugated curdlan to glass coverslips via carbonyldiimidazole (CDI)-mediated hydroxyl-amine coupling (Fig. S1A). The second kind of coverslip was a bi-functional curdlan-Pam3 substrate, which would simultaneously activate both receptors. To prepare such coverslips, we further coated curdlan-coated glass substrates with biotinylated bovine serum albumin (BSA) via physical adsorption, and then conjugated them with biotinylated Pam3 (biotin-Pam3) via biotin-streptavidin linkers (Fig. S1B). The third type of coverslip we prepared was a Pam3-only substrate, intended to activate only TLR2 receptors We coated bare glass substrates first with poly-L-lysine (PLL), then with biotin-BSA via physical adsorption, and finally with biotin-Pam3 via biotin-streptavidin linkage (Fig. S1C). In order to make direct quantitative comparisons between the behaviors of cells on the latter two substrates, it was necessary to match the surface density of Pam3 on the Pam3-only substrate with that on the bifunctional curdlan-Pam3 substrate. To do this, a PLL coating was used. The matched Pam3 density was confirmed by counting the surface density of fluorescently labeled streptavidin (linkers for biotin-Pam3) using TIRF microscopy (Fig. S2). Without PLL, the maximum surface density of biotin-Pam3 we could achieve on bare glass substrates was too low to match that on the bifunctional curdlan-Pam3 substrates. Using the special fabrication procedures described, curdlan-only and Pam3-only substrates display the same surface density of curdlan and Pam3, respectively, as the bifunctional curdlan-Pam3 substrates. In addition to the three types of coverslips with ligands, we used PLL-only glass substrates without ligands as a control (Fig. S1D).

To image TLR2 and Dectin-1 using dSTORM, cells were fixed after 10 min activation on the different substrates and then stained with antibodies to visualize the distribution of the receptors. Dectin-1 GFP was labeled with anti-GFP primary antibody and Cy3B-labeled secondary antibody, whereas TLR2 was labeled with anti-TLR2 primary antibody and Alexa Fluor 647 (AF647)-labeled secondary antibody. The localization inaccuracy of dSTORM was estimated to be 18 ± 8 nm for the Cy3B channel and 22 ± 8 nm for the AF647 channel (Fig. S3). After Gaussian rendering, images showed that both TLR2 and Dectin-1 receptors formed discrete nanoclusters on the cell membrane across the entire area where the cell contacted the substrate (Fig. 1A). Cells stimulated on ligand-coated substrates exhibited significantly larger numbers of receptor nanoclusters than resting cells on PLL-only control substrates, indicating that receptor nanoclusters form as a result of ligand stimulation.

**Figure 1.**
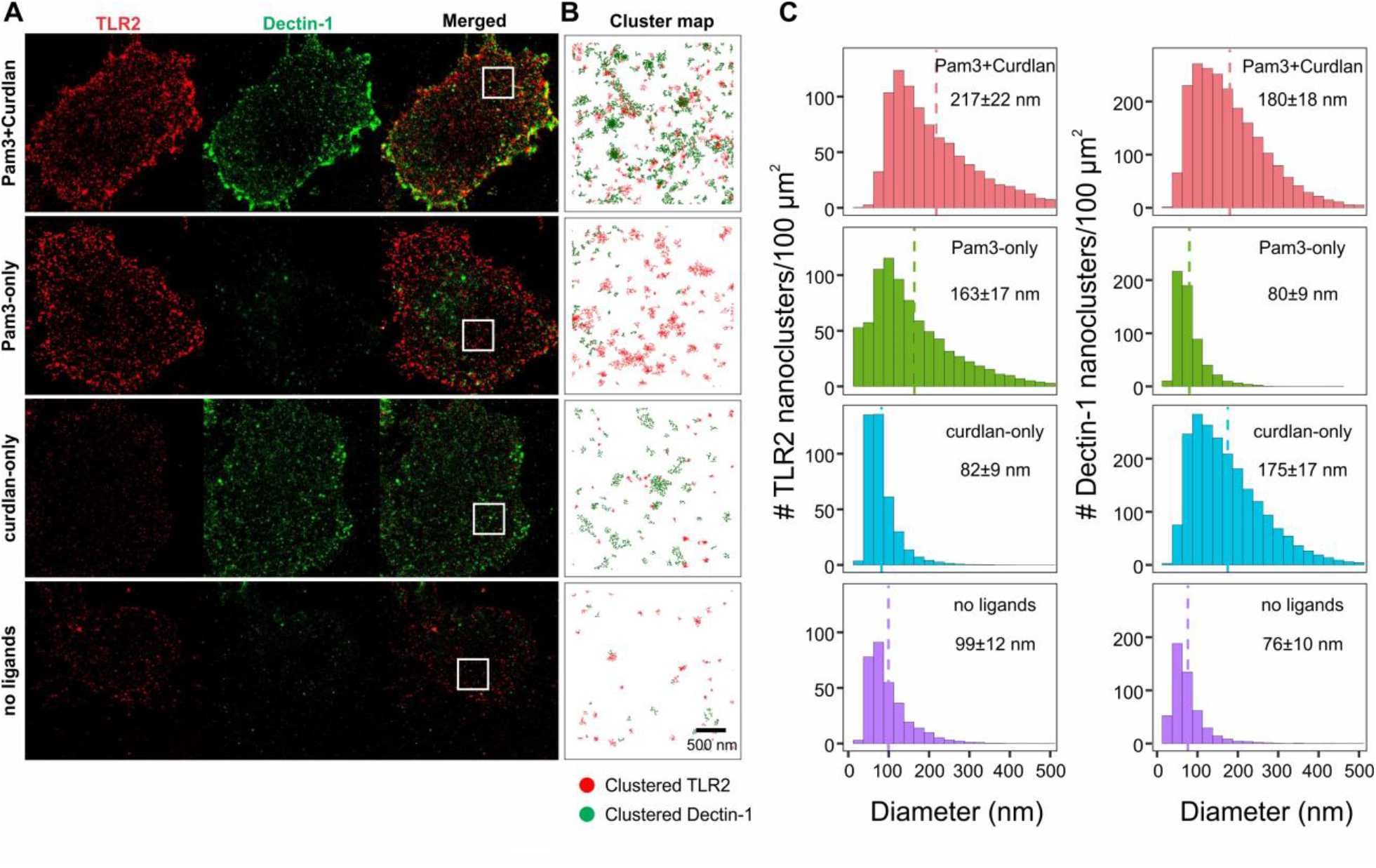
Nanoscale organization of TLR2 and Dectin-1 in macrophage cell membranes on various substrates. **(A)** Two-color dSTORM images of immunofluorescently labeled TLR2 and Dectin-1 in RAW264.7 cell membranes that were in contact with ligand-coated and control substrates. Cells were seeded on substrates for 10 min before fixation. **(B)** Cluster maps show a magnified view of the distribution of receptor nanoclusters within the areas outlined by white boxes (3 μm × 3 μm) shown in **(A)**. Receptor clusters were identified using the topological mode analysis tool (ToMATo) method. Single-molecule localizations of TLR2 (red dots) and Dectin-1 (green dots) in clusters. **(C)** Histograms showing the surface densities of TLR2 (left) and Dectin-1 (right) nanoclusters at given cluster diameters. Dotted vertical lines indicate the average cluster diameter of each type of receptors. Legend in each graph indicates mean ± SEM of N cells from 3 independent experiments. For TLR2 nanoclusters, N = 28 cells (Pam3+Curdlan), 26 cells (Pam3-only), 22 cells (curdlan-only), and 22 cells (no ligands). For Dectin-1 nanoclusters, N = 41 cells (Pam3+Curdlan), 36 cells (Pam3-only), 42 cells (curdlan-only), and 29 cells (no ligands).

In our analysis of the nanoclusters, we first performed dSTORM post-image processing to eliminate artifacts such as multi-blinking from single fluorophores. We then used a topological mode analysis tool (ToMATo) ^30^ to distinguish receptor nanoclusters (Fig. 1B) from non-clustered localizations that can result from non-clustered receptors, non-specifically adsorbed antibodies, and other factors. The ToMATo method uses a persistence-based clustering algorithm.^31^ The details of this analysis are shown in Fig. S4, and the means of validating the method, in Fig. S5. The means by which the parameters were optimized are explained in the experimental section. Through this analysis, we found a significantly higher density of Dectin-1 localizations at the interface between the cell and the substrate after stimulation by curdlan-only substrates than when exposed to the PLL control. Similarly, more TLR2 localizations per unit area were observed at the cell-substrate interface upon stimulation by Pam3-only substrates than for the PLL control (Fig. S6). This confirms previous studies that have already shown that ligand-bound phagocytic receptors, such as Dectin-1 and Fc gammar receptors (FcγRs), accumulate at the host cell-pathogen contact area.^36, 37^ Synergistic activation of both Dectin-1 and TLR2 by the bifunctional substrate resulted in an even higher density of TLR2 localizations at the cell-substrate contact area than for the unifunctional Pam3-only substrate. The number of Dectin-1 localizations, on the other hand, was unchanged from that observed for the unifunctional curdlan-only substrate (Fig. S6). It was found that 78.04 ± 7.78% of TLR2 localizations and 80.43 ± 5.26% of Dectin-1 localizations were organized in nanoclusters when a stimulating ligand was present (Fig. S7). In contrast, only 13.86 ± 2.29% of TLR2 localizations and 15.64 ± 2.45% of Dectin-1 localizations in resting cells formed nanoclusters (Fig. S7). This indicates that Dectin-1 and TLR2 are mostly randomly distributed in resting cells, and that they reorganize into nanoclusters upon ligand stimulation. Interestingly, synergistic activation of both Dectin-1 and TLR2 on the bifunctional substrate had no effect on the percentage of receptor localizations that formed nanoclusters.

From the ToMATo analysis, we also quantified the size of receptor nanoclusters. The few TLR2 nanoclusters found in resting cells were 99 ± 24 nm in diameter at a surface density of 372 ± 53 clusters/100 μm^2^. In cells exposed to Pam3 stimulation, the nanoclusters became larger (163 ± 22 nm) and more densely distributed (803 ± 64 nanoclusters/100 μm^2^) (Fig. 1C and Fig. S8). Similarly, the Dectin-1 clusters exposed to curdlan stimulation were 175±30 nm in diameter at a higher density of 1415±101 nanoclusters/100 μm^2^, compared to a mean diameter of 76 ± 18 nm at an average density of 312 ± 71 nanoclusters/100 μm^2^ in resting cells (Fig. 1C and Fig. S8). We showed that the formation of the observed receptor nanoclusters, in both resting and activated cells, requires an intact actin cytoskeleton. In control experiments (Fig. S9), cells were seeded on the different substrates for 5 min and then incubated for another 5 min with 2.5 μg/ml cytochalasin D (cytoD); a substance which disrupts actin. The surface density of TLR2 nanoclusters decreased to ∼ 8-41 nanoclusters/100 μm^2^ and their sizes decreased to ∼ 32-39 nm, regardless of the ligands presented on the substrate. Similarly, the surface density of Dectin-1 nanoclusters decreased to ∼ 7-52 nanoclusters/100 μm^2^ and their sizes decreased to ∼ 37-72 nm.

Interestingly, synergistic activation of TLR2 and Dectin-1 resulted in larger TLR2 clusters (217 ± 38 nm) than did non-synergistic activation, but had little effect on the size of Dectin-1 clusters (180 ± 32 nm). The activation of TLR2 had no effect on the surface density of activated Dectin-1 clusters, nor did the activation of Dectin-1 affect the surface density of TLR2 clusters (Fig. S8). This result, combined with the observation of TLR2 localizations suggest that Dectin-1 activation enhances the formation of TLR2 nanoclusters, but TLR2 activation has no effect on the formation Dectin-1 nanoclusters. This may be related to the different functions of Dectin-1 and TLR2 in phagocytosis. Unlike TLR2, Dectin-1 is responsible for triggering actin polymerization to mediate the engulfment of phagocytic targets.^36, 38^ It is possible that the actin polymerization triggered by Dectin-1 activation stabilizes TLR2 nanoclusters and promotes the formation of more clusters.

### 3.2. Partial overlap between nanoclusters of TLR2 and Dectin-1 upon simultaneous co-activation

An intriguing observation from the ToMATo analysis is that TLR2 and Dectin-1 clusters do not completely colocalize even during synergistic activation (Fig. 1B and Fig. 2A). We determined that, upon synergistic activation, the centroid-to-centroid nearest neighbor distance between pairs consisting of a Dectin-1 cluster and a TLR2 cluster is 109 ± 14 nm (Fig. S10). By comparison, the combined radii of TLR2 and Dectin-1 nanoclusters is notably larger; 193 ± 35 nm. This suggests that the Dectin-1 clusters and TLR2 clusters partially overlap during their synergistic activation. To quantitatively test whether the nanoclusters of TLR2 and Dectin-1 were indeed partially overlapped, we quantified the degree of overlap by adapting a coordinate-based colocalization analysis.^33^ This method was previously used for quantifying receptor colocalization in T cells^34^ and macrophage cells.^39^ In this analysis, the density gradient of single-molecule colocalizations between two types of receptors is correlated and the correlation coefficient is used as the degree of colocalization (DoC) score for each localized molecule (Fig. 2B and S11). The DoC scores range from −1 (perfectly segregated) through 0 (uncorrelated) to +1 (perfectly colocalized), and a threshold of DoC score ≥0.4 was used in previous studies to indicate colocalization between two single molecule localizations.^34, 35^ Based on this, we then calculated the degree of overlap between nanoclusters, defined as the percentage of each species of single-molecule localizations that had DoC scores ≥0.4 (Fig. 2C). By this definition, the degree of overlap of Dectin-1 clusters with TLR2 is equivalent to the fraction of surface area of Dectin-1 clusters that overlap with TLR2 clusters. The same applies to the degree of overlap of TLR2 clusters with Dectin-1.

**Figure 2.**
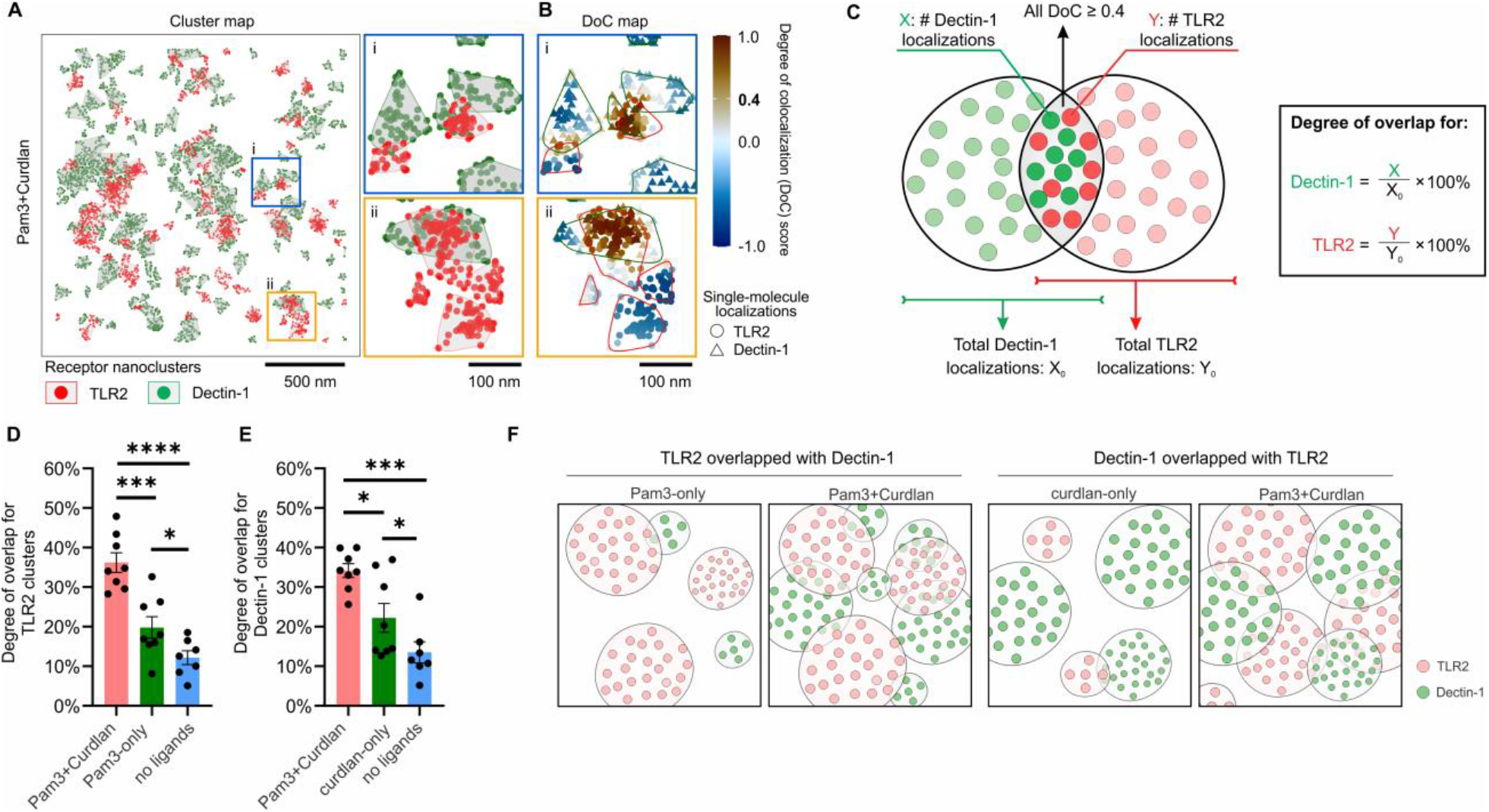
Quantification of spatial overlap between TLR2 and Dectin-1 nanoclusters. **(A)** Cluster maps of TLR2 and Dectin-1 in RAW264.7 cell membranes in contact with the curdlan-Pam3 substrates. Cells were seeded for 10 min before fixation. Single-molecule localizations of TLR2 (red dots) and Dectin-1 (green dots) are indicated. **(B)** Degree of colocalization (DoC) analysis of TLR2 and Dectin-1 single-molecule localizations within the magnified areas (boxes i and ii) indicated in (**A)**. DoC scores ranging from -1 to 1 are color-coded and assigned to each localization, with -1 indicating segregation, 0 indicating non-colocalization, and 1 indicating complete overlap. **(C)** Schematic illustration demonstrating the definition of the degree of overlap between nanoclusters. **(D-E)** Quantification of the degree of overlap for TLR2 clusters overlapped with Dectin-1 (**D**) and that for Dectin-1 clusters overlapped with TLR2 (**E**). Data are presented as mean ± SEM of N cells from 2 independent experiments. N= 8 cells from 5 images (Pam3+Curdlan), 8 cells from 5 images (Pam3-only), 8 cells from 5 images (curdlan-only), and 7 cells from 5 images cells (no ligands). Statistical significance is highlighted by p values as follows: ****p <= 0.0001; **p <= 0.01; *p <= 0.05; ns p>0.05. (**F**) Schematic illustration shows that the increased number and size of receptor nanoclusters upon synergistic activation of Dectin-1 and TLR2 can lead to overlap between TLR2 and Dectin-1 nanoclusters.

To optimize the parameters, we used to analyze cluster overlap, we first determined the validity of the DoC ≥ 0.4 threshold by using simulated patterns of receptor nanoclusters with varying degrees of overlap (Fig. S12, for details see the experimental section). We optimized parameters, including the DoC threshold and the search radius, radius step, and step count, of the coordinate-based colocalization analysis, by ensuring that 96.12% of Dectin-1 localizations and 96.18% of TLR2 localizations in resting cells were identified as non-colocalized (DoC score <0.4) (Fig. S13). This is because these two receptors are expected not to colocalize on the surfaces of resting cells.^21^ Following a previously reported procedure,^34^ we further confirmed that the setting of those parameters is not affected by variation in cluster size, or the density of single molecule localizations inside clusters (Fig. S14 and S15). Finally, we validated the analysis of overlap, first, by confirming that localizations of TLR2 or Dectin-1 in visibly separate nanoclusters from dSTORM images had negative DoC scores (Fig. S16). Secondly, we showed that single localizations of receptors that were not in clusters also had negative DoC scores (Fig. S13). In contrast, single localizations of TLR2 and Dectin-1 within the visibly overlapped region between nanoclusters had DoC > 0.4 (Fig. 2 A-B and Fig. S16).

After optimizing and validating the parameters, we quantified the degree of overlap of TLR2 clusters with Dectin-1 clusters and vice versa by calculating the percentage of single localizations of DoC ≥ 0.4 within each nanocluster. The degree of overlap between TLR2 and Dectin-1 nanoclusters was ∼ 10 % in resting cells (Fig. 2D and E), which serves as a negative control. When only TLR2 was activated, the degree of overlap of TLR2 clusters with Dectin-1 was 19.80 ± 2.72%. Likewise, when only Dectin-1 was activated, the degree of overlap of Dectin-1 clusters with TLR2 was 22.43 ± 3.27%. Upon synergistic activation of TLR2 and Dectin-1, the degree of overlap increased further to 36.21±2.47% for TLR2 clusters overlapped with Dectin-1 and 34.17±1.75% for Dectin-1 clusters overlapped with TLR2 (Fig. 2D and E). We have shown in Fig. 1 that more receptor clusters are formed after stimulation by a specific ligand. The surface density of TLR2 clusters increased 2.2-fold after Pam3 stimulation and that of Dectin-1 clusters increased 4.5 fold. Moreover, the TLR2 clusters increased in size by 1.6-fold after synergistic activation of both TLR2 and Dectin-1. It is very likely that the increased degree of overlap between TLR2 and Dectin-1 clusters, compared to when only one receptor is activated, is due to the increased number of receptor clusters and the larger size of TLR2 clusters (Fig. 2F).

### 3.3. Effect of TLR2 and Dectin-1 crosstalk on receptor activation and signaling at single receptor clusters

We next investigated how the synergistic activation of TLR2 and Dectin-1 affects their signaling competency in single receptor clusters. To determine when TLR2 was activated, we used the recruitment of the cytoplasmic adaptor protein, myeloid differentiation primary response 88 (MyD88) as an indicator.^40^ Activation of Dectin-1 was indicated by the phosphorylation of recruited spleen tyrosine kinase (Syk). After Dectin-1 is activated, its cytoplasmic motifs are phosphorylated and become docking sites for Syk. Once recruited to the motifs, Syk is activated and phosphorylated to mediate downstream signaling.^41^ Just as in all our dSTORM sample preparation, we fixed cells after 10 min activation on the different substrates, and immunostained the samples for TLR2 and MyD88, or Dectin-1 and phosphor-Syk (pSyk). As expected for activated receptors, many of the TLR2 and Dectin-1 clusters colocalized with puncta of MyD88 and pSyk, respectively, as revealed in dSTORM images and cluster maps from the ToMATo analysis (Fig. 3A). The level of immunostained MyD88 and pSyk was significantly less in resting cells (Fig. S17 and S18).

**Figure 3.**
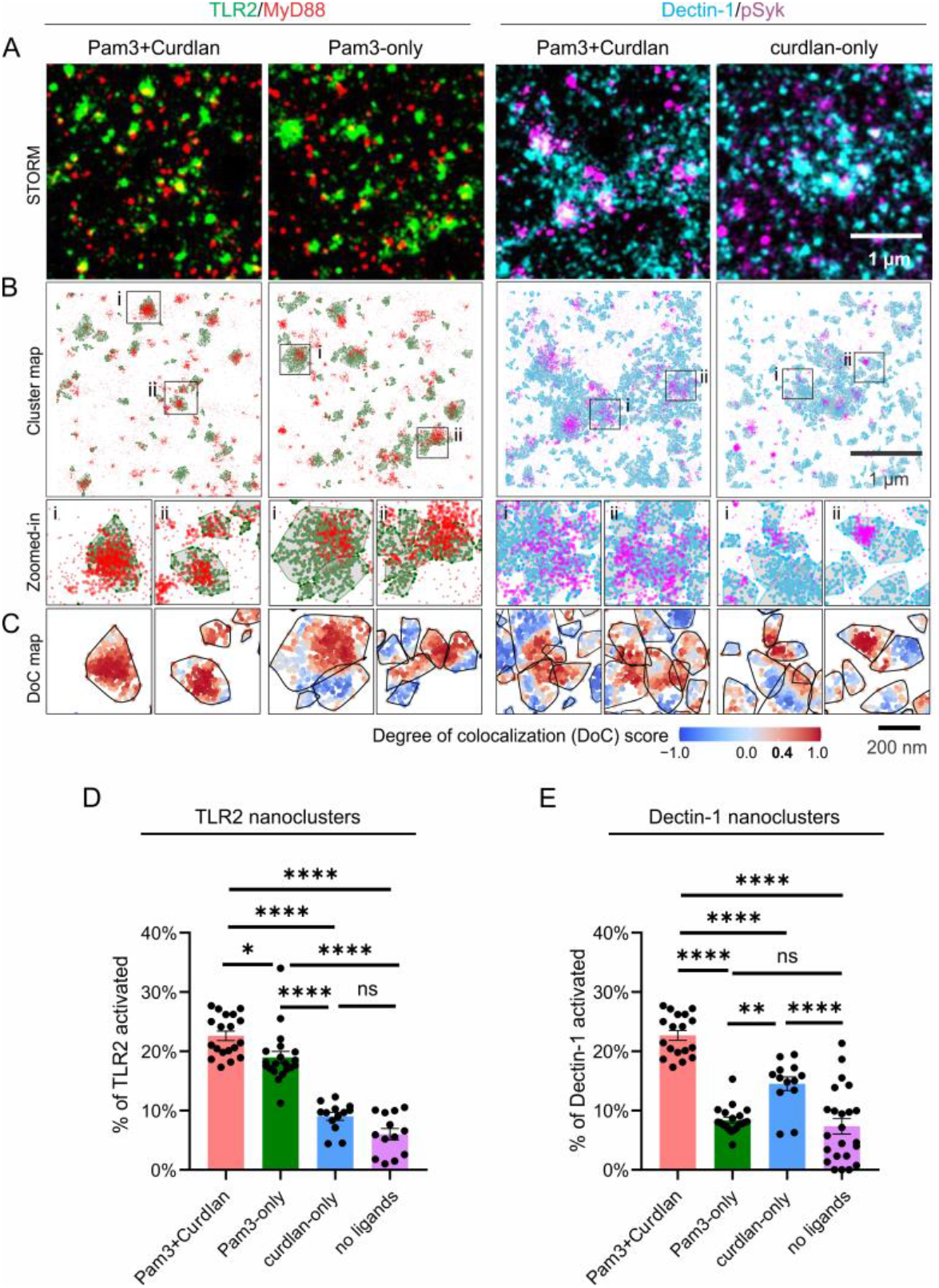
Quantification of the activation of TLR2 and Decitn-1 nanoclusters. **(A)** Representative two-color dSTORM images of immunofluorescently labeled TLR2 and MyD88, and of immunofluorescently labeled Dectin-1 and pSyk. All cells were seeded on ligand-functionalized glass coverslips for 10 min before fixation. **(B)** Cluster maps of TLR2 (green) and Dectin-1 (cyan) overlapped with the single-molecule localizations of MyD88 (red) and pSyk (magenta), respectively, corresponding to Raw dSTORM images in **(A)**. TLR2 and Dectin-1 localizations that were identified as in nanoclusters are enclosed in polygons. **(C)** Upper portion: Degree of colocalization (DoC) analysis of TLR2 and MyD88, and of Dectin-1 and pSyk in areas outlined by boxes i and ii in Panel **B**. Lower portion: Activation of TLR2 or Dectin-1 is indicated by the color-coded DoC scores between TLR2 and MyD88 or Dectin-1 and pSyk, respectively. Receptor nanocluster contours are highlighted with black lines. **(D-E)** Quantification of the percentage of activated TLR2 in nanoclusters (**D**) and perecentage of activated Dectin-1 in nanoclusters (**E**) under different stimulation conditions as indicated. Data are represented as mean ± SEM of N cells from 2 independent experiments. For TLR2 and MyD88 imaging, N=19 cells from 8 images (Pam3+Curdlan), 20 cells from 8 images (Pam3-only), 13 cells from 8 images (curdlan-only) and 13 cells from 6 images (no ligands). For Dectin-1 and pSyk imaging, N=18 cells from 7 images (Pam3+Curdlan), 20 cells from 11 images (Pam3-only), 13 cells from 8 images (curdlan) and 23 cells from 7 images (no ligands). Statistical significance is highlighted by p values as follows: ****p <= 0.0001; **p <= 0.01; *p <= 0.05; ns p>0.05.

We determined the percentage of receptors that were activated by quantifying the degree of overlap of TLR2 clusters with the activation indicator MyD88, and that of Dectin-1 clusters with pSyk (Fig. 3C). We used the same coordinate-based colocalization analysis mentioned above. The colocalization of TLR2 with MyD88 indicates its activation, and the same is true for Dectin-1 and pSyk. This is why the degree of overlap is a measure of the percentage of the receptors that were activated. Results from resting cells indicate that the basal level of activation was 7.36 ± 1.30% for Dectin-1 and 6.00 ± 0.96% for TLR2. When only Dectin-1 was activated on curdlan-only substrates, 14.52 ± 1.51% of Dectin-1 in clusters were activated and TLR2 activation was unaffected. Similarly, when only TLR2 was activated on Pam3-only substrates, 18.98 ± 1.02% of TLR2 in clusters were activated, whereas Dectin-1 was not activated. However, upon synergistic activation of both TLR2 and Dectin-1, the percentages of activation increased to 22.70 ± 0.83% for Dectin-1 and 22.61 ± 0.79% for TLR2. This demonstrates that during synergistic crosstalk, the activation of TLR2 increases the probability of Dectin-1 activation and the activation of Dectin-1 similarly enhances TLR2 activation.

Synergistic crosstalk increases the percentage of activated Dectin-1 and TLR2 during exposure to the bifunctional substrate. We sought to assess the consequences of this for intracellular immune signaling. To do this, we quantified the phosphotyrosine (pY) intensity at the level of single receptor clusters. Tyrosine phosphorylation is involved in signaling pathways deriving from both TLR2 and Dectin-1 activation. Its intensity thus serves as a general indicator of receptor signaling activities.^42-44^ Cells were exposed to 10 min. of stimulation on the various substrates of our experiment, and then fixed and immunofluorescently stained for pY. TIRF microscopy images showed the immunostained pY as discrete puncta near the cell membrane at the adhesion interface between the cell and thesubstrate (Fig. 4A). As expected, the number and fluorescence intensity of pY puncta were much higher upon activation of either or both receptors than that in resting cells. We quantified the fluorescence intensity per pY puncta using a single-particle tracking algorithm, as well as the average pY intensity per 100 µm^2^ to indicate the whole-cell level signaling intensity (Fig. 4B and C). Results from both quantifications are consistent: the pY level was similar when only Dectin-1 or only TLR2 was activated, but increased significantly upon synergistic activation of both receptors. The average pY intensity per cell was also increased by synergistic activation. This result confirms that the increased percentage of activated Dectin-1 and TLR2 receptors, caused by their synergistic crosstalk (Fig. 3), leads to increased cell signaling.

**Figure 4.**
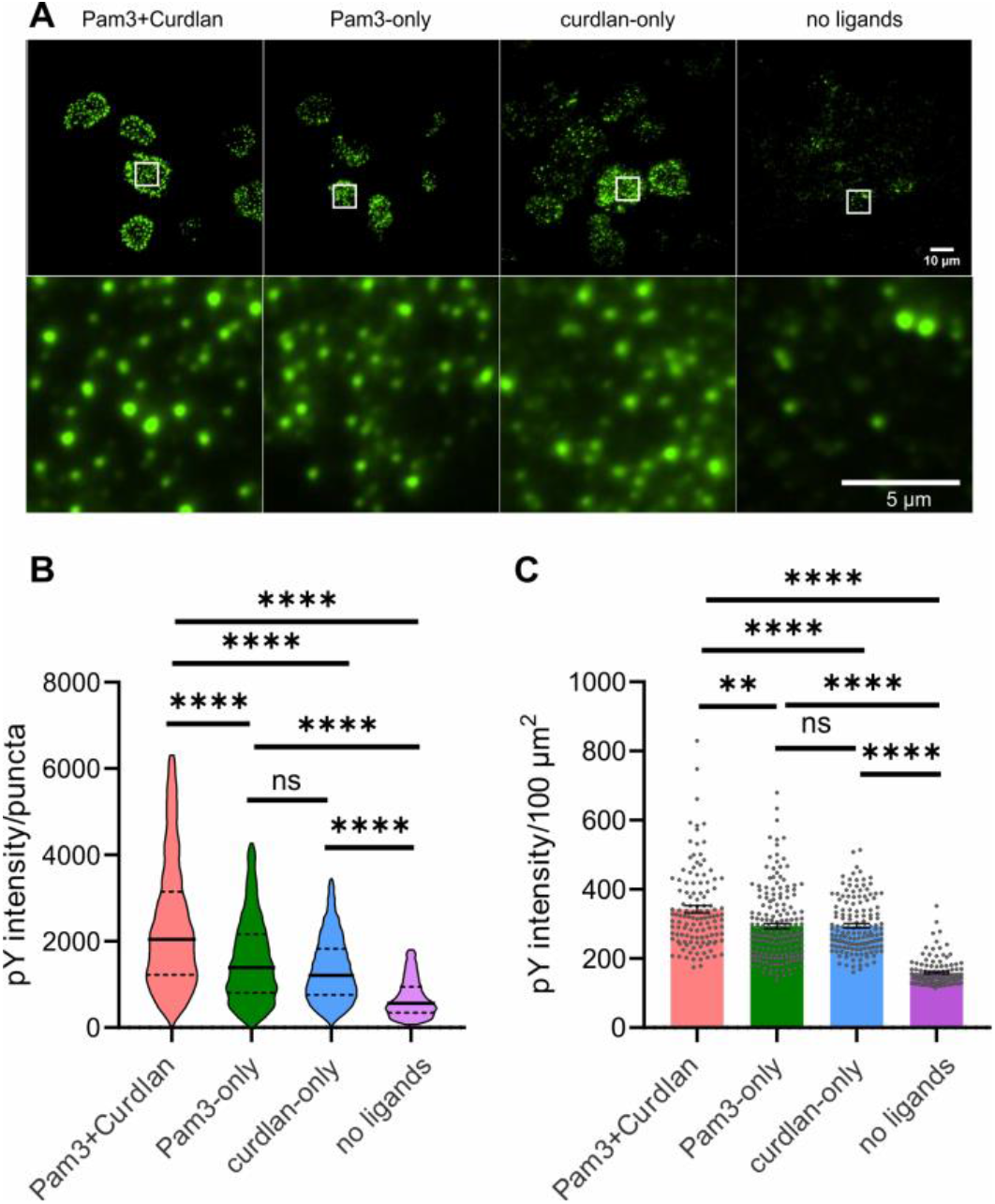
Effect of receptor activation on phosphotyrosine (pY) signaling. **(A)** Fluorescence images showing immunostained pY in RAW264.7 macrophage cells after 10 min of incubation on different substrates. Zoom-in images show pY puncta in regions of interest outlined by white boxes. **(B)** Quantification of fluorescence intensity of individual pY puncta. Data were from 2 independent experiments and presented as violin plots with median and interquartile range. N=11231 puncta from 178 images (Pam3+Curdlan), 8618 puncta from 123 images (Pam3-only), 7764 puncta from 145 images (curdlan-only) and 853 puncta from 117 images (no ligands). **(C)** Quantification of pY fluorescence intensity of individual cells. For quantitative comparison, pY intensity of each cell was obtained as intensity per cell area. Data are presented as mean ± SEM obtained from 178 cells (Pam3+Curdlan), 123 cells (Pam3-only), 145 cells (curdlan-only), and 117 cells (no ligands). Statistical significance is highlighted by p values as follows: ****p <= 0.0001; **p <= 0.01; *p <= 0.05; ns p>0.05.

## 4. Conclusions

Dectin-1 and TLR2 function synergistically to elicit innate immune responses,^13-18^ but how their signaling crosstalk occurs on the receptor cluster level has not been investigated. In this study, we revealed, with super-resolution dSTORM, the nanoscale organization of TLR2 and Dectin-1 in the plasma membrane of RAW264.7 macrophage cells. We characterized the consequences of this receptor organization for intracellular immune signaling. Our results demonstrate that, after ligand stimulation, Dectin-1 and TLR2 separately form discrete nanoclusters that are 217 ± 22 nm and 180 ± 18 nm in diameter. The sizes of nanoclusters were shown to be much smaller than what we reported previously using diffraction-limited TIRF imaging,^23, 45^ We found that Dectin-1 and TLR2 nanoclusters do not colocalize but are partially overlapped during their signaling crosstalk (schematic in Fig. 5). Compared to when only one type of receptor is activated, the synergistic activation of Dectin-1 and TLR2 leads to a larger fraction of activated receptors and consequently to stronger cell signaling as indicated by the level of tyrosine phosphorylation. Our results support a model in which Dectin-1 and TLR2 achieve synergistic signaling crosstalk through interaction at the interface between the receptor clusters.

**Figure 5.**
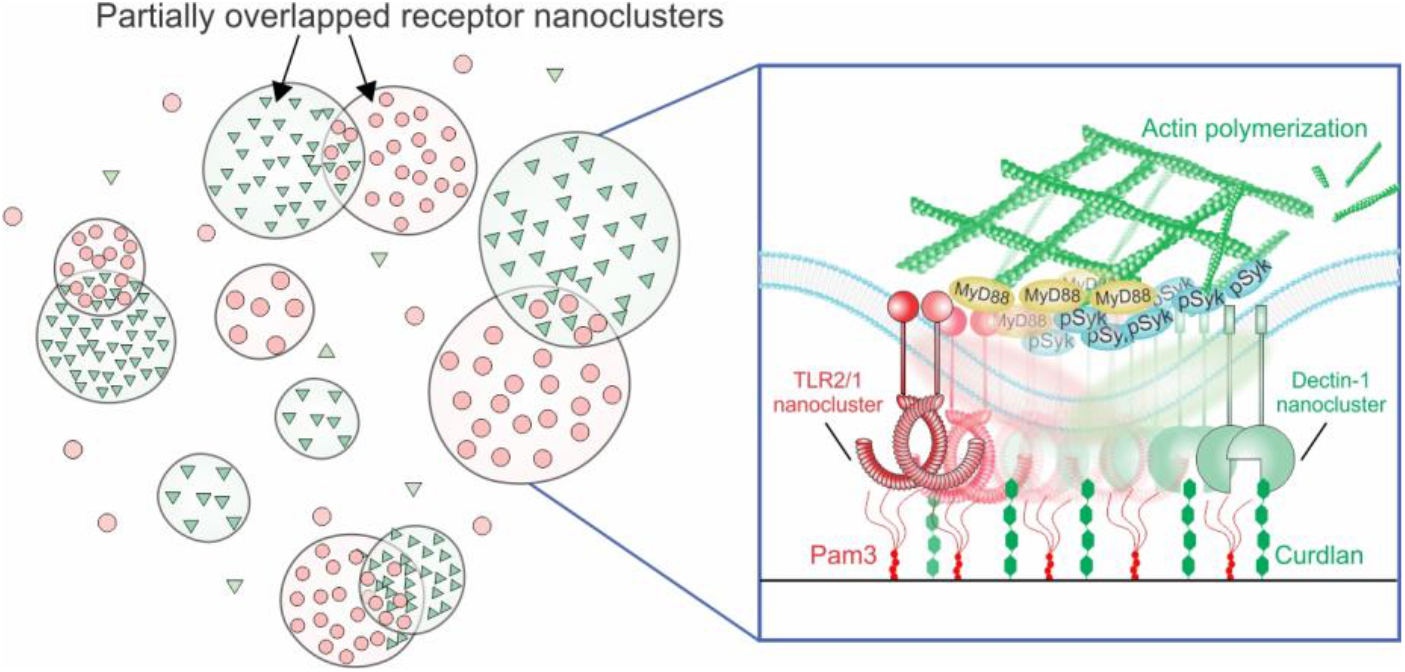
Schematic illustration showing the nanoscale organization of TLR2 and Dectin-1 receptors at macrophage surface upon synergistic activation. Upon synergistic stimulation, TLR2 and Dectin-1 receptors organize in partially overlapped nanoclusters. The overlapping interfaces between nanoclusters allow receptors and signaling proteins to interact for signaling crosstalk. Actin polymerization supports the formation of TLR2 and Dectin-1 nanoclusters and their synergistic crosstalk.

Many previous studies have suggested that Dectin-1 and TLR2 colocalize to function synergistically. Our previous study challenged this view by showing that Dectin-1 and TLR2 form discrete clusters that reside within nanoscale proximity during signaling crosstalk.^23^ In this study, we demonstrated that the receptor clusters are, in fact, partially overlapped. With dSTORM imaging and a coordinate-based colocalization analysis, we not only directly visualized the overlap between the Dectin-1 and TLR2 nanoclusters, but also quantified the degree of overlap. We do not yet know how TLR2 and Dectin-1 interact within the overlapped region to coordinate their synergistic crosstalk. We speculate that the overlapped interfaces between TLR2 and Dectin-1 nanoclusters serve as signaling “hotspots”, where both types of receptors can intermix and directly interact with one another, as well as exchanging proximal signaling molecules that converge their signals. For instance, studies suggested that TLR2 and Dectin-1 signals converge on caspase recruitment domain family member 9 (CARD9) to activate the transcription factor NF-κB.^46-50^ When receptor clusters interact at the interfaces, different types of receptors may be able to intermix or separate more rapidly to regulate their crosstalk, compared to the colocalization models in which different species of receptors completely intermix. Interestingly, we have also found recently that TLR2 and FcγR form discrete receptor nanoclusters that become partially overlapped during synergistic signaling.^51^ It remains to be explored further whether this model of receptor clusters intermixing at interfaces is a common mechanism shared by many immune receptors that have signaling crosstalk.

## Supporting material

Supporting material includes fig. S1-S18 and can be found online.

## Acknowledgments

We are indebted to Prof. Bin Dong for helping us build the STORM microscope. Research reported in this publication was supported by the National Institute of General Medical Sciences of the National Institutes of Health under Award Number R35GM124918. The content is solely the responsibility of the authors and does not necessarily represent the official views of the National Institutes of Health.

## Author Contributions

M. L. and Y. Y. designed project; M. L., C. V. and M. B performed the STORM imaging and other experiments; M. L. analyzed data; and M. L. and Y. Y. wrote the paper.

