## Supplemental figures for "Spatial organization of Dectin-1 and TLR2 during synergistic crosstalk revealed by super-resolution imaging"

**This PDF file includes:**

Figures. S1-S18

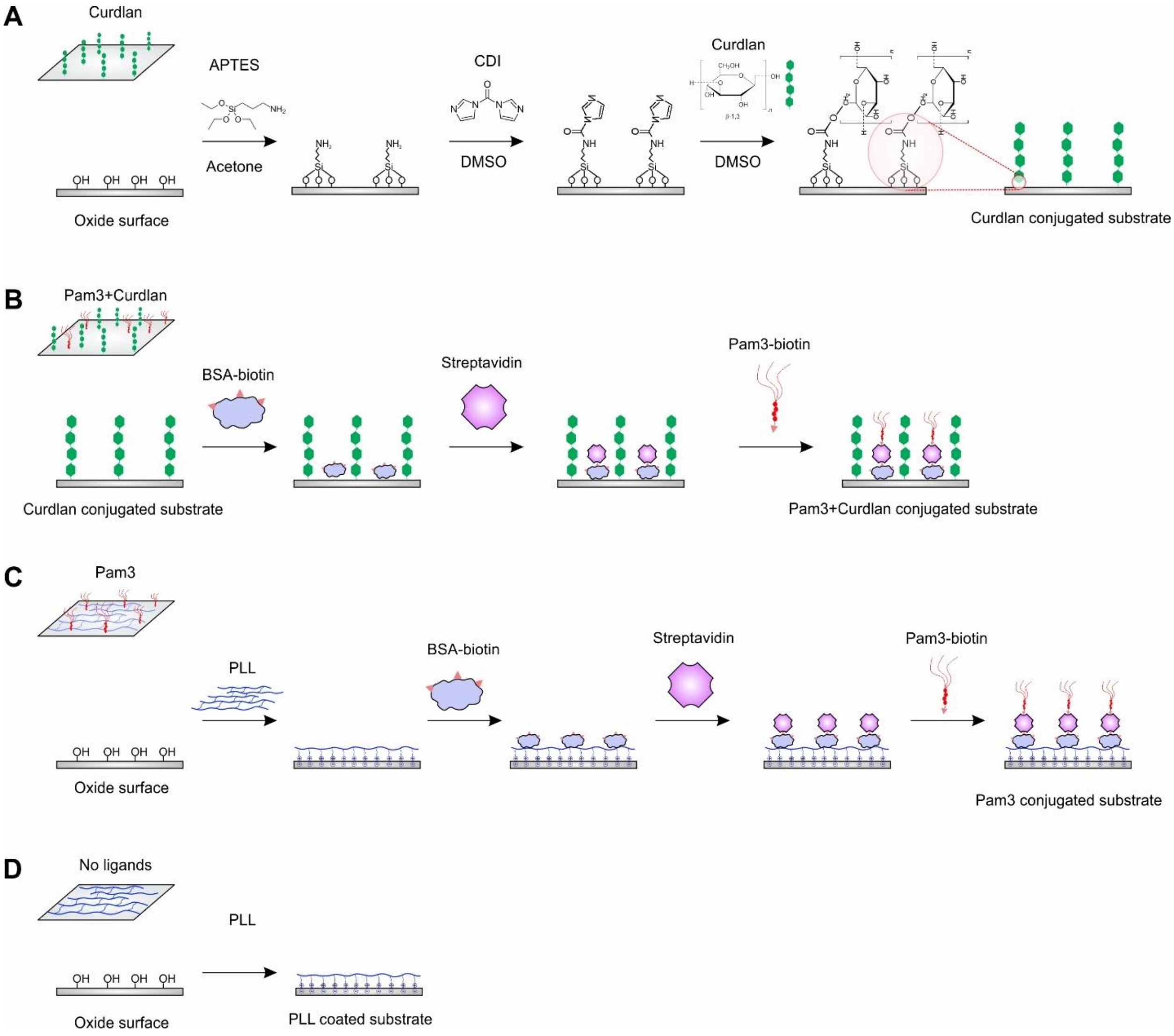


**Fig. S1: Schematic illustration of the fabrication procedures for ligand functionalized glass coverslips. (A)** Pam3-only surface. **(B)** Curdlan-only surface. **(C)** Pam3 and curdlan bi-functionalized surface. **(D)** poly-L-lysine (PLL)-coated surface.

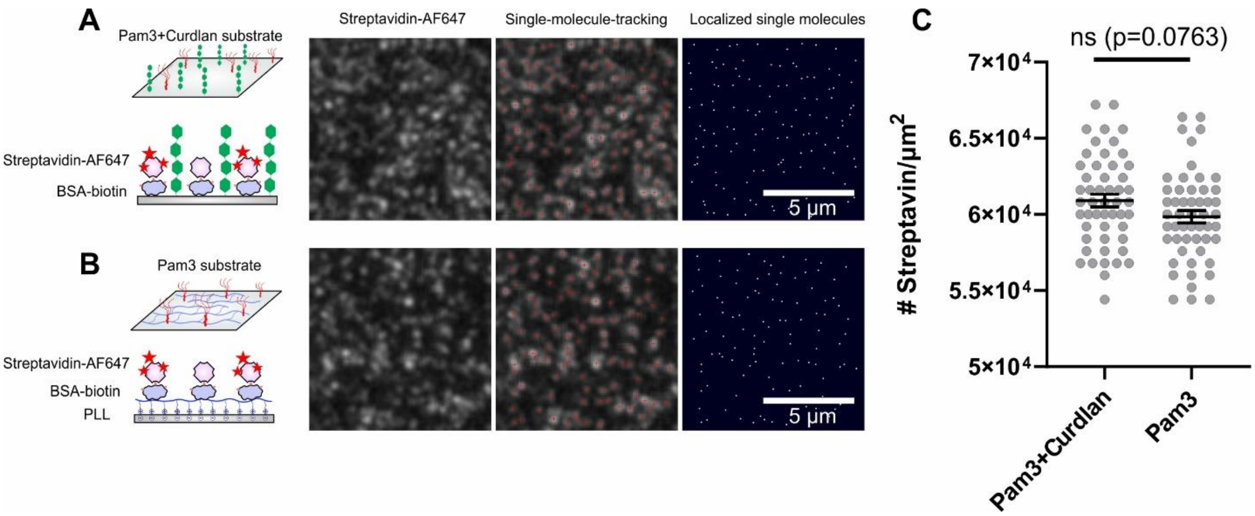


**Fig. S2: Quantification of Pam3 density on glass coverslips. (A-B)** Representative TIRF images and single-molecule localizations of the Alexa Fluor 647 (AF647) labeled streptavidin on **(A)** the curdlan-only substrate and **(B)** the PLL-coated substrate. AF647 labeled streptavidin was mixed with unlabeled streptavidin at a molar ratio of 1: 2000 for single-molecule visualization. **(C)** Quantification of the surface density of streptavidin on the curdlan-Pam3 substrate and the Pam3-only substrate. Each point represents the result from one image. Each data set of the scatter plot represents mean ± SEM obtained from a total of N = 51 images (Pam3+Curdlan) and 51 images (Pam3), respectively, from 3 independent measurements each. Statistical significance, assessed using the Mann-Whitney U test, is highlighted by P values as follows: ns $P > 0.05$.

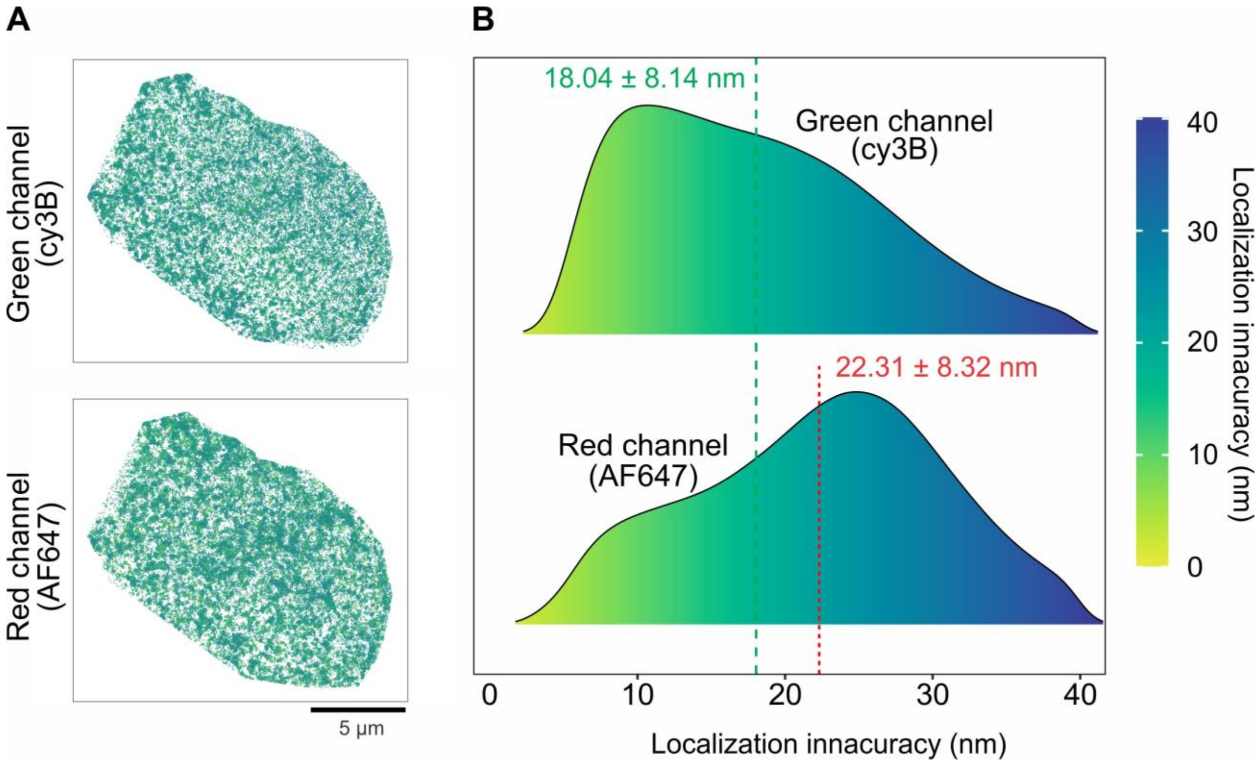


**Fig. S3: Quantification of localization inaccuracy of dSTORM images. (A)** Representative heat maps showing the single-molecule localizations of Cy3B-labeled Dectin-1 (green channel) and AF647-labeled TLR2 (red channel) in a single cell. Each localization is color-coded according to its localization inaccuracy. **(B)** Probability distributions of localization inaccuracy of the green channel and the red channel. Data in each graph were from 10 cells from 3 independent experiments. Legend in each graph indicates mean ± SEM.

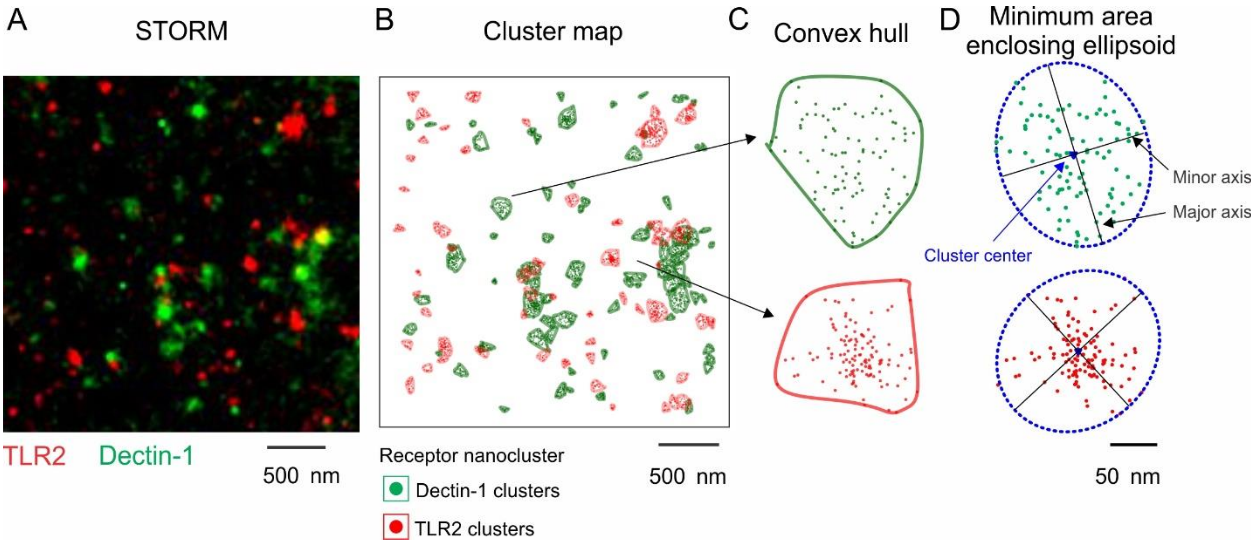


**Fig. S4: Identification of receptor nanoclusters from dSTORM images. (A)** A representative dSTORM image of immunostained TLR2 (red) and Dectin-1 (green) in the plasma membrane of RAW264.7 macrophage cells seeded on the curdlan-Pam3 substrate. **(B)** Single-molecule localizations of TLR2 and Dectin-1 in **(A)** were analyzed using a topological mode analysis tool (ToMATo) to identify nanoclusters. Receptor molecules in nanoclusters are enclosed in polygons. **(C)** The convex hull of the clustered single localizations shows the representative nanoclusters of Dectin-1 (green) and TLR2 (red). **(D)** The diameter of a nanocluster was calculated using the mean value of the major and minor axes of the bounding ellipse. The bounding ellipse was identified using the minimal area enclosing all localizations within a nanocluster.

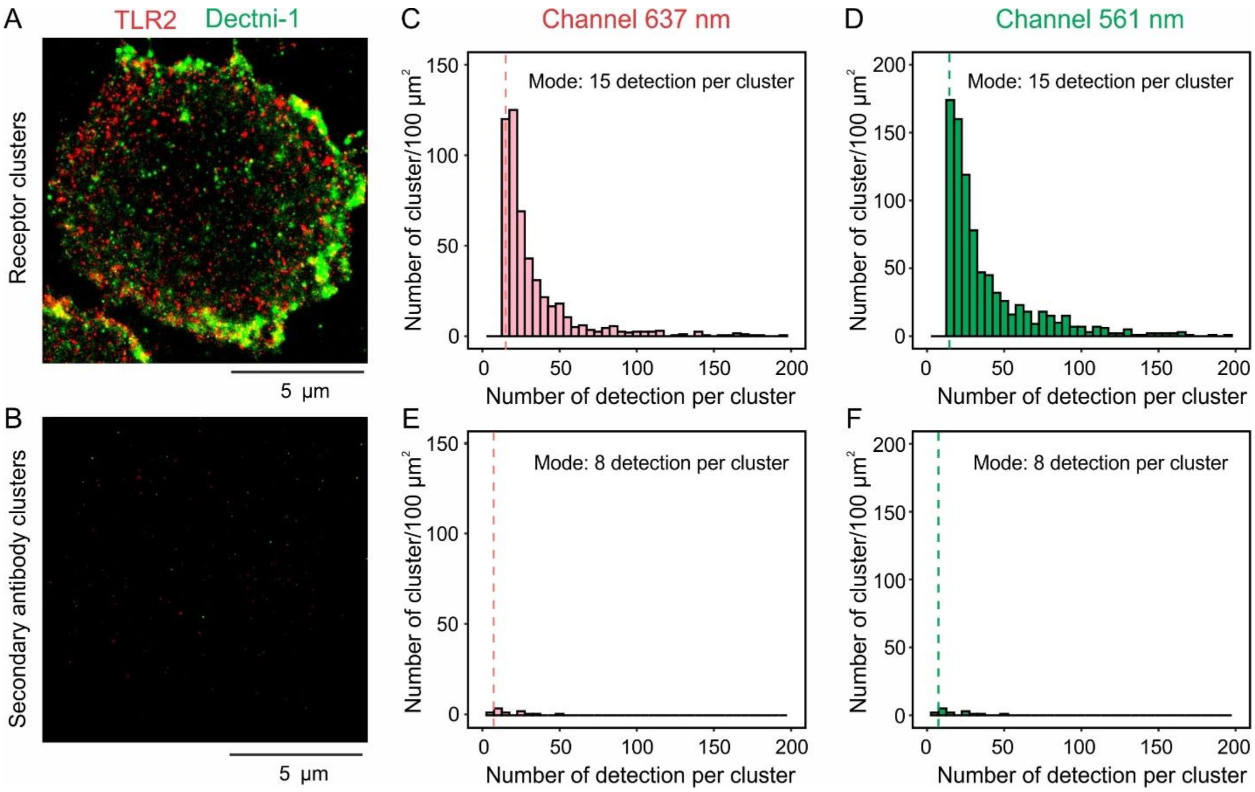


**Fig. S5: Quantification of the number of localizations in receptor nanoclusters and antibody nanoclusters. (A)** A representative dSTORM image of immunofluorescently labeled TLR2 and Dectin-1 nanoclusters in the membrane of a RAW264.7 macrophage cell that was activated on a bi-functional curdlan-Pam3 substrate for 10 min. **(B)** A representative dSTORM image of fluorescent secondary antibodies that were non-specifically adsorbed to a cell sample on the curdlan-Pam3 substrate. **(C-D)** Histograms show the surface density of TLR2 (**C**) and Dectin-1 nanoclusters (**D**) versus the number of single-molecule localizations detected in each receptor nanocluster. **(E-F)** Histograms show the surface density of AF647-labeded secondary antibody nanoclusters (**E**) and Cy3B-labeled secondary antibody nanoclusters (**F**) non-specifically formed at the cell surface versus the number of single-molecule localizations detected in each antibody nanocluster. Vertical lines indicate the mode value of the number of detected single-molecule localizations per nanocluster. Each data set was obtained from 15 non-overlapping regions (3 μm × 3 μm) in 5 cells from 2 independent samples.

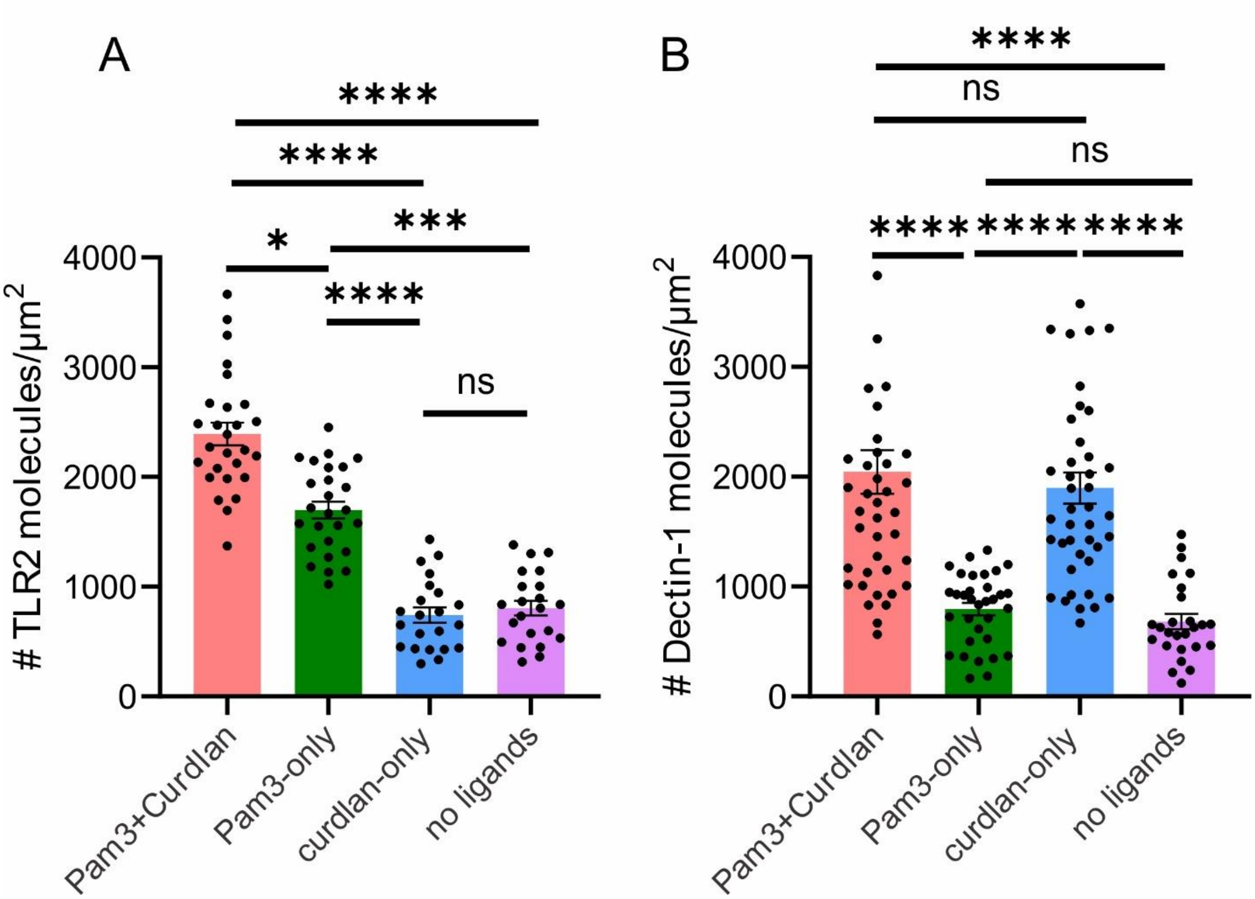


**Fig. S6: Quantification of the surface density of receptor molecules. (A, B)** The surface density of TLR2 and Dectin-1 in cells on different substrates as indicated. Each point represents the result from one cell. Horizontal bars in the bar graphs represent mean ± SEM of N cells from 3 independent experiments. For TLR2 nanoclusters, N = 27 (Pam3+Curdlan), 26 (Pam3-only), 22 (curdlan-only), and 22 cells (no ligands). For Dectin-1 nanoclusters, N = 41 (Pam3+Curdlan), 33 (Pam3-only), 42 (curdlan-only), and 26 cells (no ligands). Statistical significance is highlighted by P values as follows: ****p <= 0.0001; **p <= 0.01; *p <= 0.05; ns p>0.05.

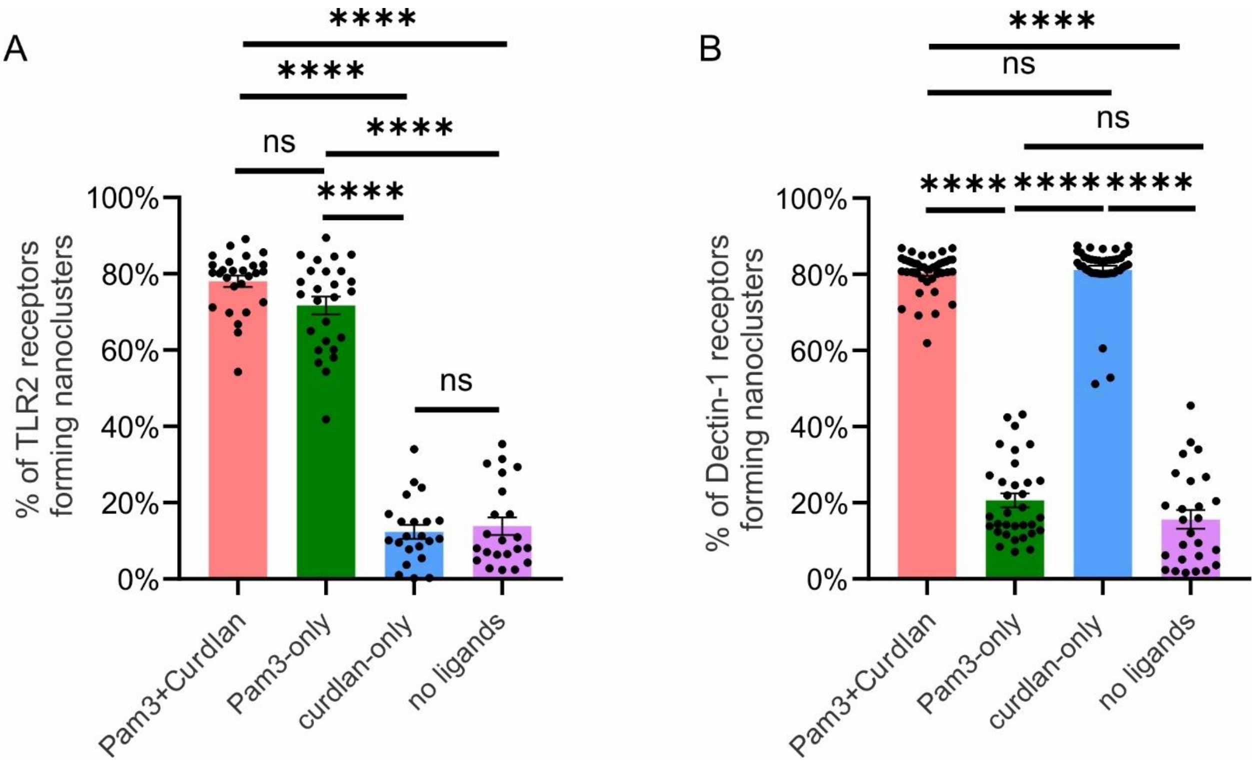


**Fig. S7: Quantification of the percentage of receptor molecules that form nanoclusters. (A, B)** The percentage of TLR2 molecules and Dectin-1 molecules that form nanoclusters in cells on different substrates. Each data point is result from a cell. Horizontal bars in the bar graphs represent mean ± SEM of N cells from 3 independent experiments. For TLR2 nanoclusters, N = 27 (Pam3+Curdlan), 26 (Pam3-only), 22 (curdlan-only), and 22 cells (no ligands). For Dectin-1 nanoclusters, N = 41 (Pam3+Curdlan), 33 (Pam3-only), 42 (curdlan-only), and 26 cells (no ligands). Statistical significance is highlighted by P values as follows: ****p <= 0.0001; **p <= 0.01; *p <= 0.05; ns p>0.05.

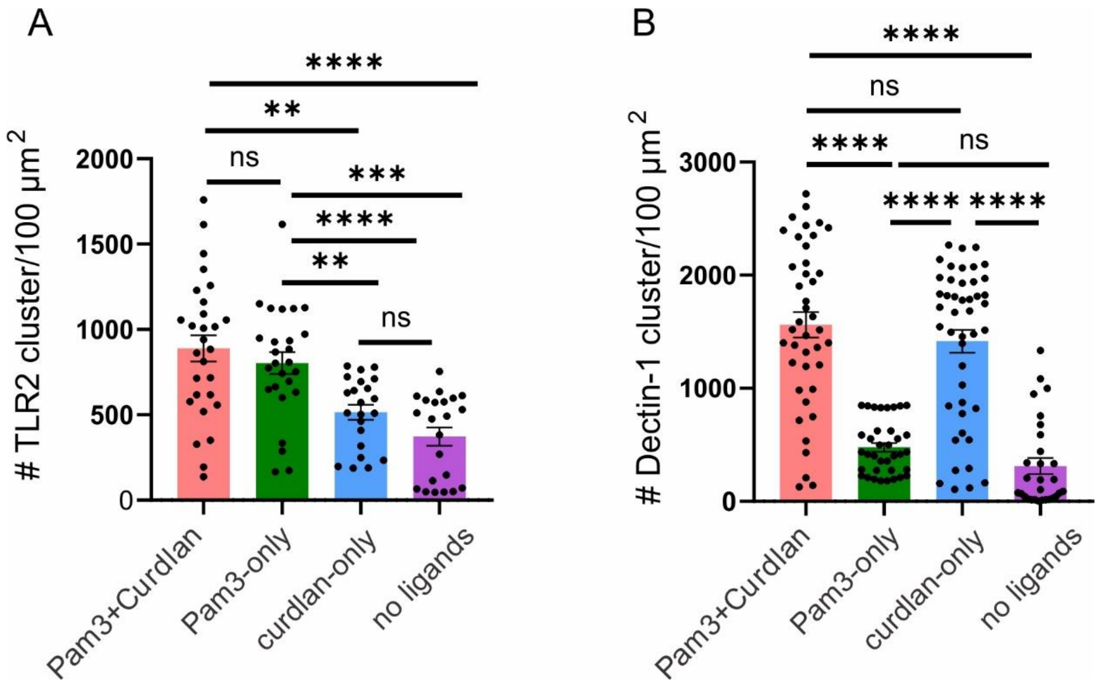


**Fig. S8: Quantification of the surface density of receptor nanoclusters. (A, B)** The surface density of TLR2 nanoclusters and Dectin-1 nanoclusters in the membrane of cells on different substrates. Each point represents the result from one cell. Horizontal bars in the bar graphs represent mean ± SEM of N cells from 3 independent experiments. For TLR2 nanoclusters, N=28 (Pam3+Curdlan), 26 (Pam3-only), 22 (curdlan), and 22 cells (no ligands). For Dectin-1 nanoclusters, N=41 (Pam3+Curdlan), 36 (Pam3-only), 45 (curdlan-only), and 29 cells (no ligands). Statistical significance is highlighted by P values as follows: ****p <= 0.0001; **p <= 0.01; *p <= 0.05; ns p>0.05.

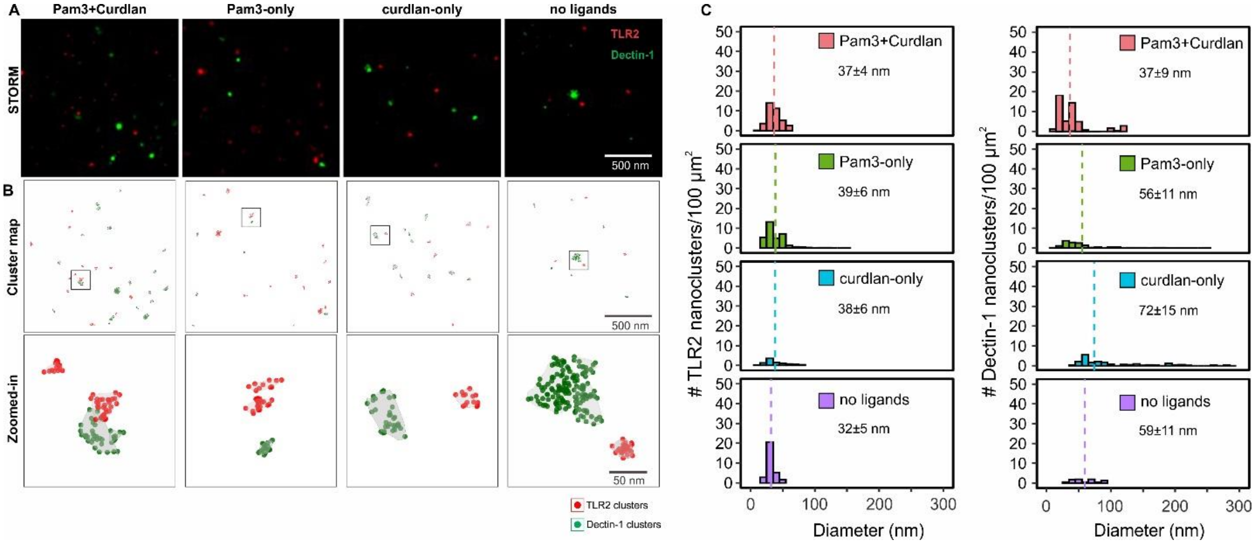


**Fig. S9: Effect of actin cytoskeleton on the formation of TLR2 and Dectin-1 nanoclusters. (A)** Two-color dSTORM images of immunofluorescently labeled TLR2 and Dectin-1 in the membrane of RAW264.7 cells on different substrates as indicated. Cells were seeded on substrates for 5 min and then incubated with cytochalasin D (cytoD) for 5 min before fixation. **(B)** Cluster maps show the distribution of receptor nanoclusters. The receptor clusters were identified using the topological mode analysis tool (ToMATo) method. Single-molecule localizations of TLR2 and Dectin-1 in clusters are shown as red dots and green dots, respectively. **(C)** Histograms show the surface density of TLR2 and Dectin-1 nanoclusters at given cluster diameters. Dotted vertical lines indicate the average cluster diameter of each type of receptors. Legend in each graph indicates mean ± SEM of N cells from 2 independent experiments. N = 13 (Pam3+Curdlan), 8 (Pam3-only), 11 (curdlan-only), and 8 cells (no ligands).

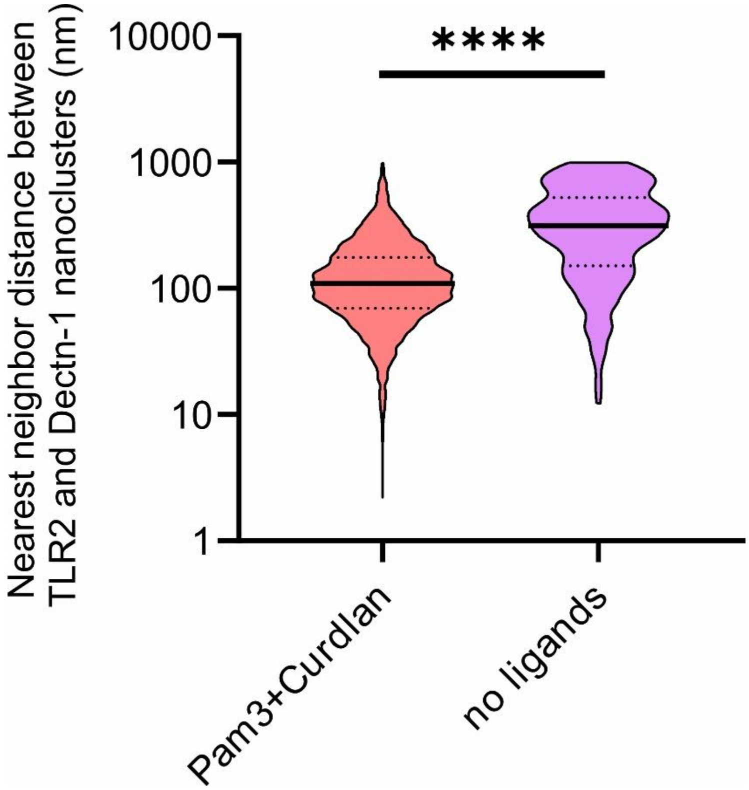


**Fig. S10: Quantification of nearest neighbor distance between TLR2 and Dectin-1 nanoclusters.** Data are presented as violin plots with the median and interquartile range from N pairs of TLR2 and Dectin-1 nanoclusters collected from 2 independent experiments. N = 6923 nanocluster pairs from 5 images in 8 cells (Pam3+Curdlan), and N = 384 nanocluster pairs from 5 images in 7 cells pairs (no ligands). Statistical significance is highlighted by p values as follows: ****p <= 0.0001; **p <= 0.01; *p <= 0.05; ns p>0.05.

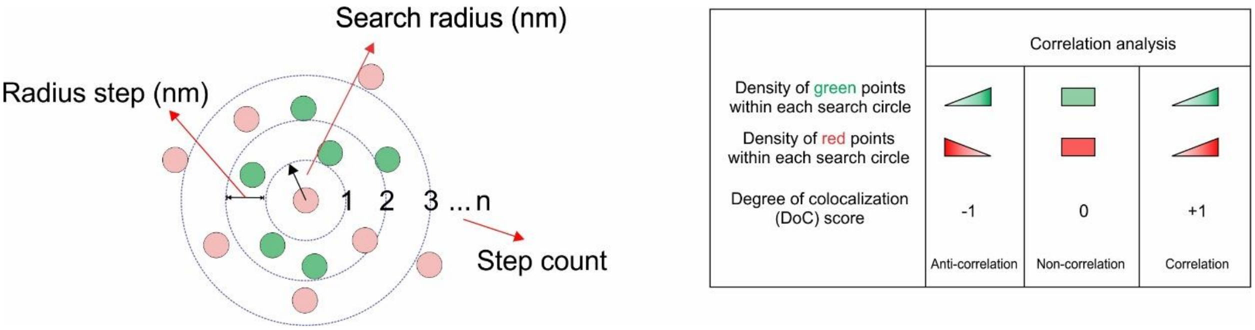


**Fig. S11: Coordinate-based colocalization (CBC) analysis and parameter determination.** The local density (number of localizations per unit area) of each channel (red and green) is calculated at increasing search radius sizes with a fixed radius step and a maximum step count. This generates density gradients of both channels. The correlation coefficient of two gradients represents a degree of colocalization (DoC) score for each localized molecule. DoC scores range from -1 to 1, with -1 indicating anti-correlation (perfect segregation), 0 for non-correlation (non-colocalization), and 1 for perfect correlation (perfect colocalization).

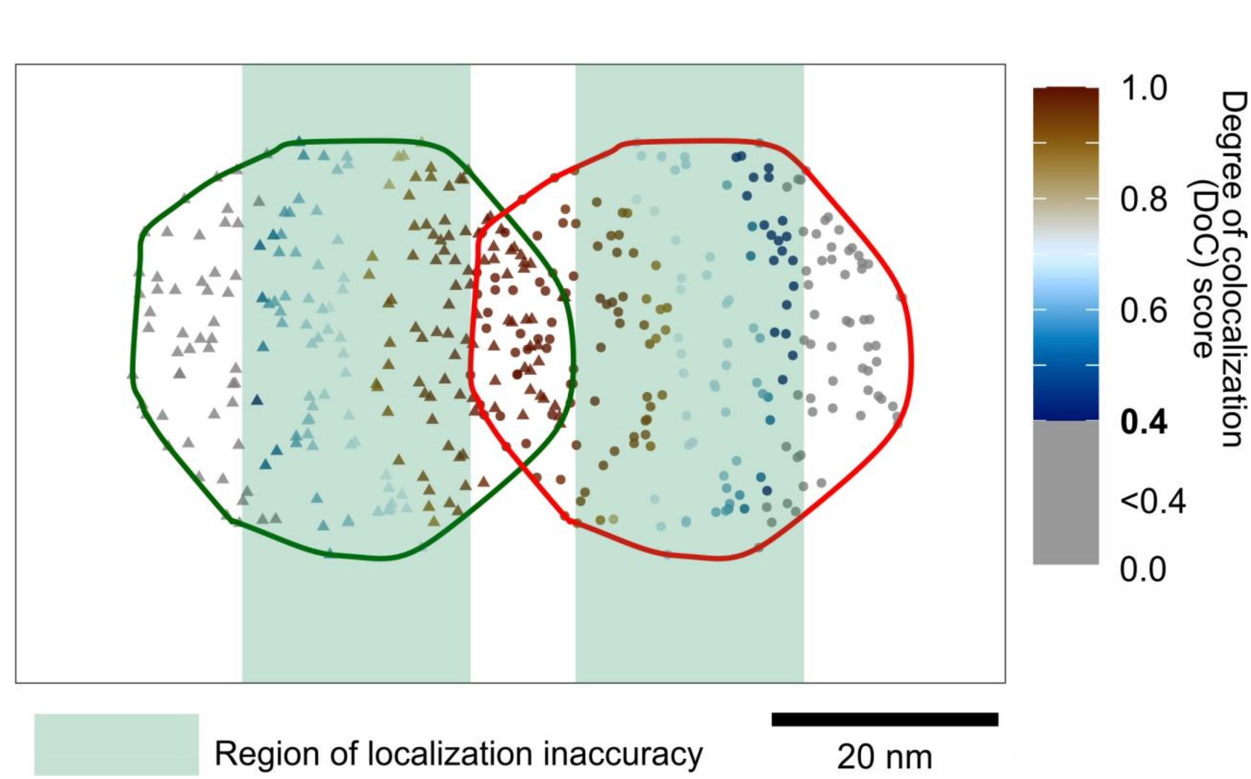


**Fig. S12: Simulation for DoC score threshold identification.** Two identical receptor nanoclusters (35 nm in diameter) were laterally translated for a distance of 30 nm. CBC analysis was performed using a search radius of 10 nm, a radius step of 10 nm, and a step count of 25 to calculate the DoC scores at each single molecule localization. The color-coded points represent localizations with DoC scores ≥0.4. Green shaded areas represent the regions of localization inaccuracy of the dSTORM imaging (i.e. the error of the measurement).

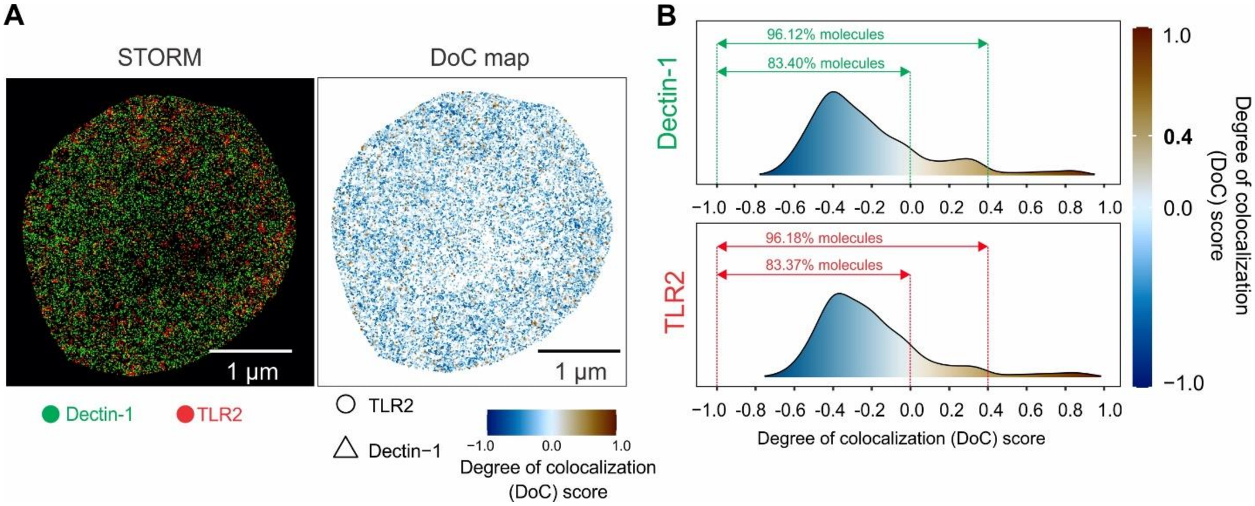


**Fig. S13: Quantification of DoC score for non-clustered receptors. (A)** A representative dSTROM image of non-clustered TLR2 and Dectin-1 in resting cells (left) and the DoC color-coded score map (right). The Doc score was calculated at each single-molecule localization using a search radius of 10 nm, a radius step of 10 nm, and a step count of 25. **(B)** DoC score distributions of non-clustered Dectin-1 and non-clustered TLR2. Each data set was from 9 cells from 3 independent experiments.

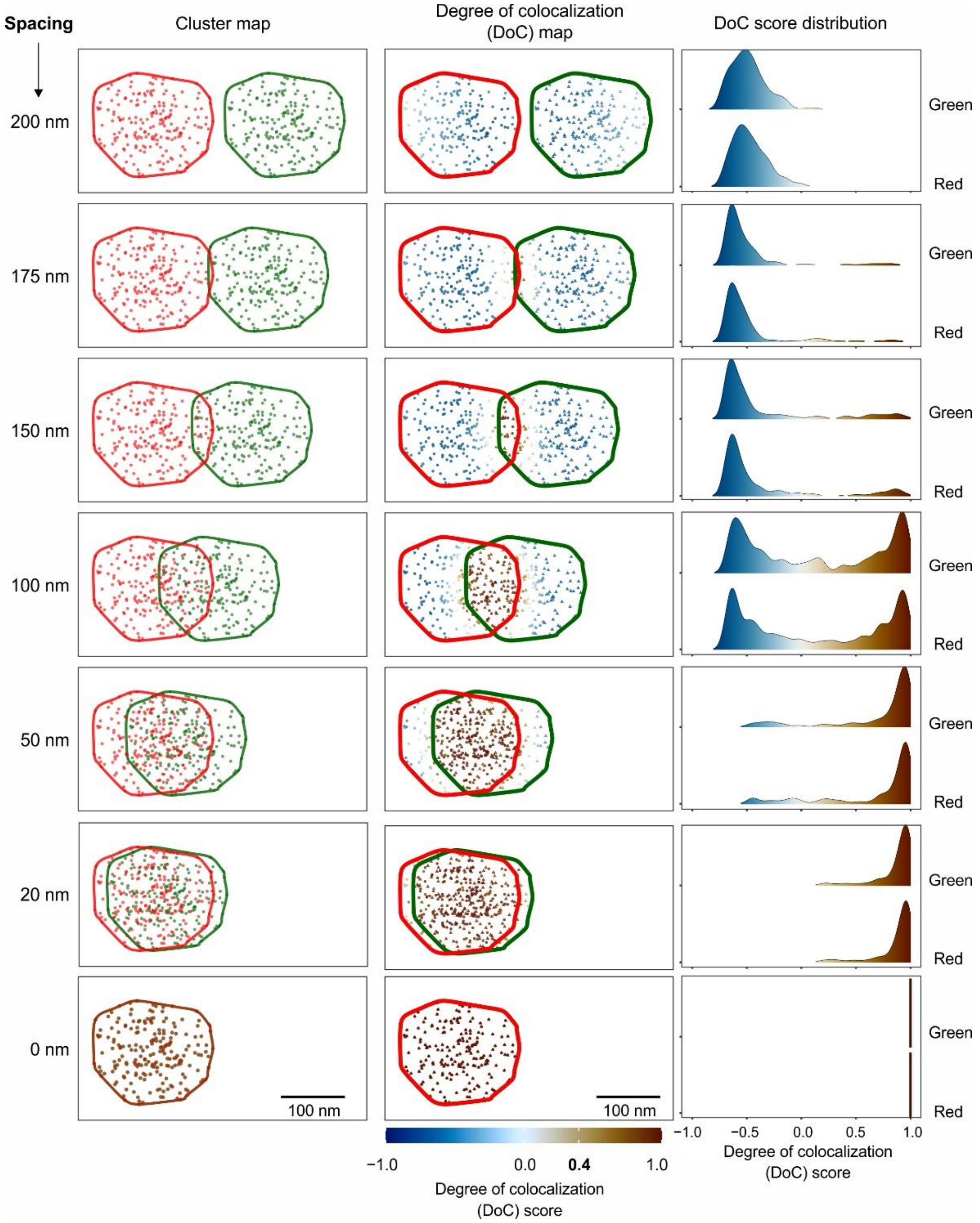


**Fig. S14: Validation of CBC analysis with low-density nanoclusters.** Two identical receptor nanoclusters (182 nm in diameter and 206 localizations) are laterally translated for different distances from 0 to 200 nm as indicated. CBC analysis was performed using a search radius of 10 nm, a radius step of 10 nm and a step count of 25 to calculate the DoC scores (color-coded) at each localization.

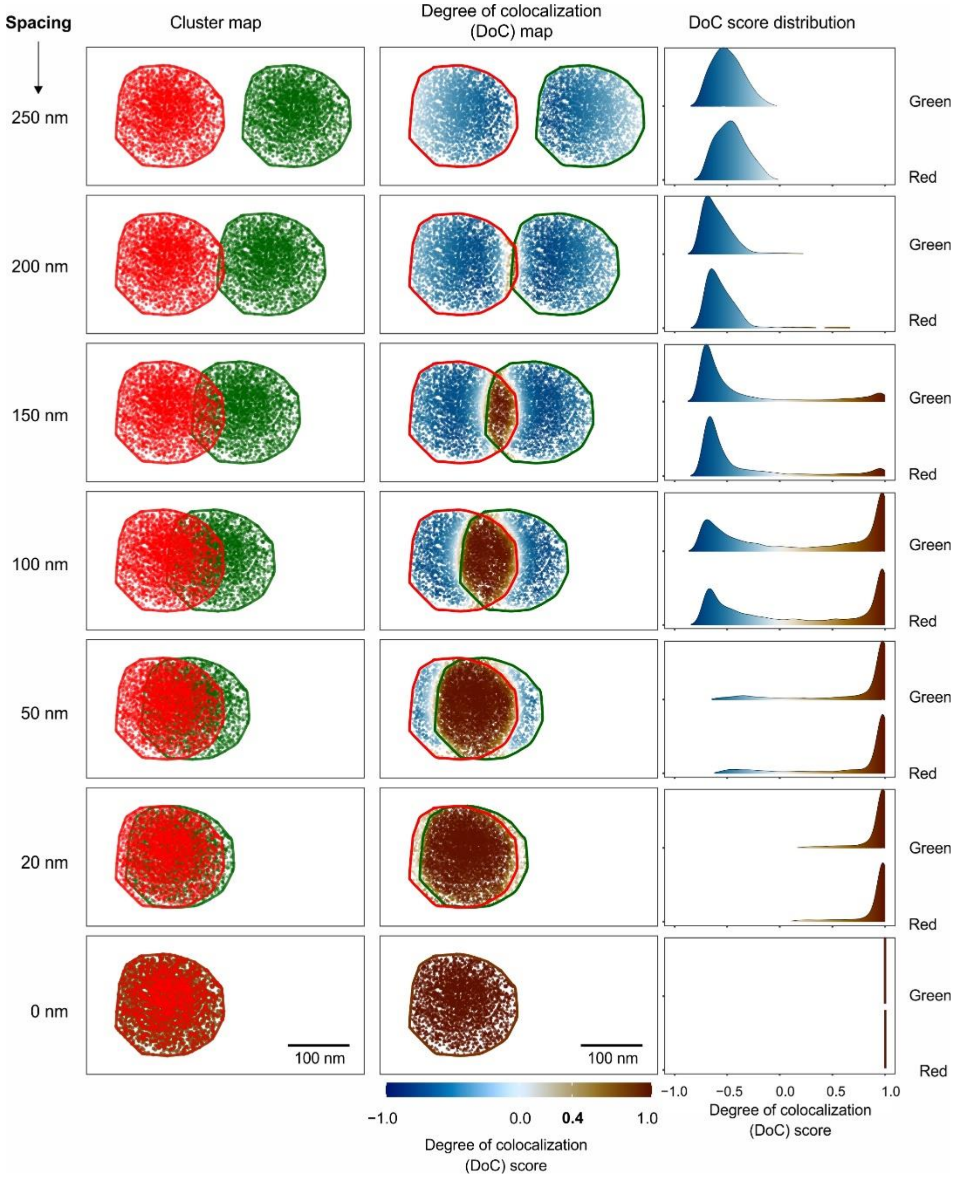


**Fig. S15: Validation of CBC analysis with high-density nanoclusters.** Two identical receptor nanoclusters (218 nm in diameter and 2927 localizations) are laterally translated for various distances from 0 to 250 nm as indicated. CBC analysis was performed using a search radius of 10 nm, a radius step of 10 nm and a step count of 25 to calculate the DoC scores (color-coded) at each localization.

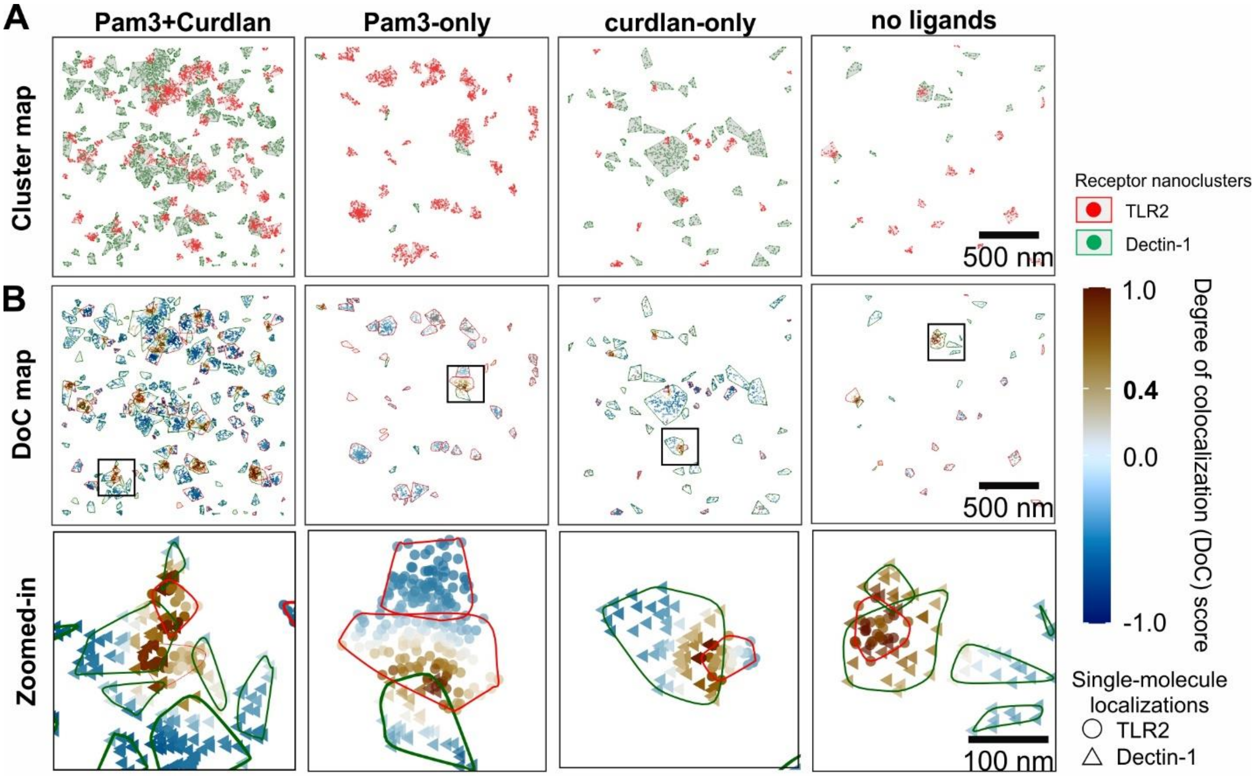


**Fig. S16: Quantification of the degree of overlap between TLR2 and Dectin-1 nanoclusters. (A)** Representative cluster maps of TLR2 and Dectin-1 receptors in the membrane of RAW264.7 cells on different substrates. Single-molecule localizations of TLR2 and Dectin-1 are shown as red and green dots, respectively. Localizations within nanoclusters are enclosed in polygons. **(B)** Degree of colocalization (DoC) analysis of TLR2 and Dectin-1 single-molecule localizations shown in **(A)**. DoC scores ranging from -1 to 1 are color-coded and assigned to each localization, with -1 indicating segregation, 0 indicating non-colocalization and 1 indicating colocalization. Zoomed regions in black boxes show the DoC maps of representative TLR2 and Dectin-1 nanoclusters that partially overlap with each other.

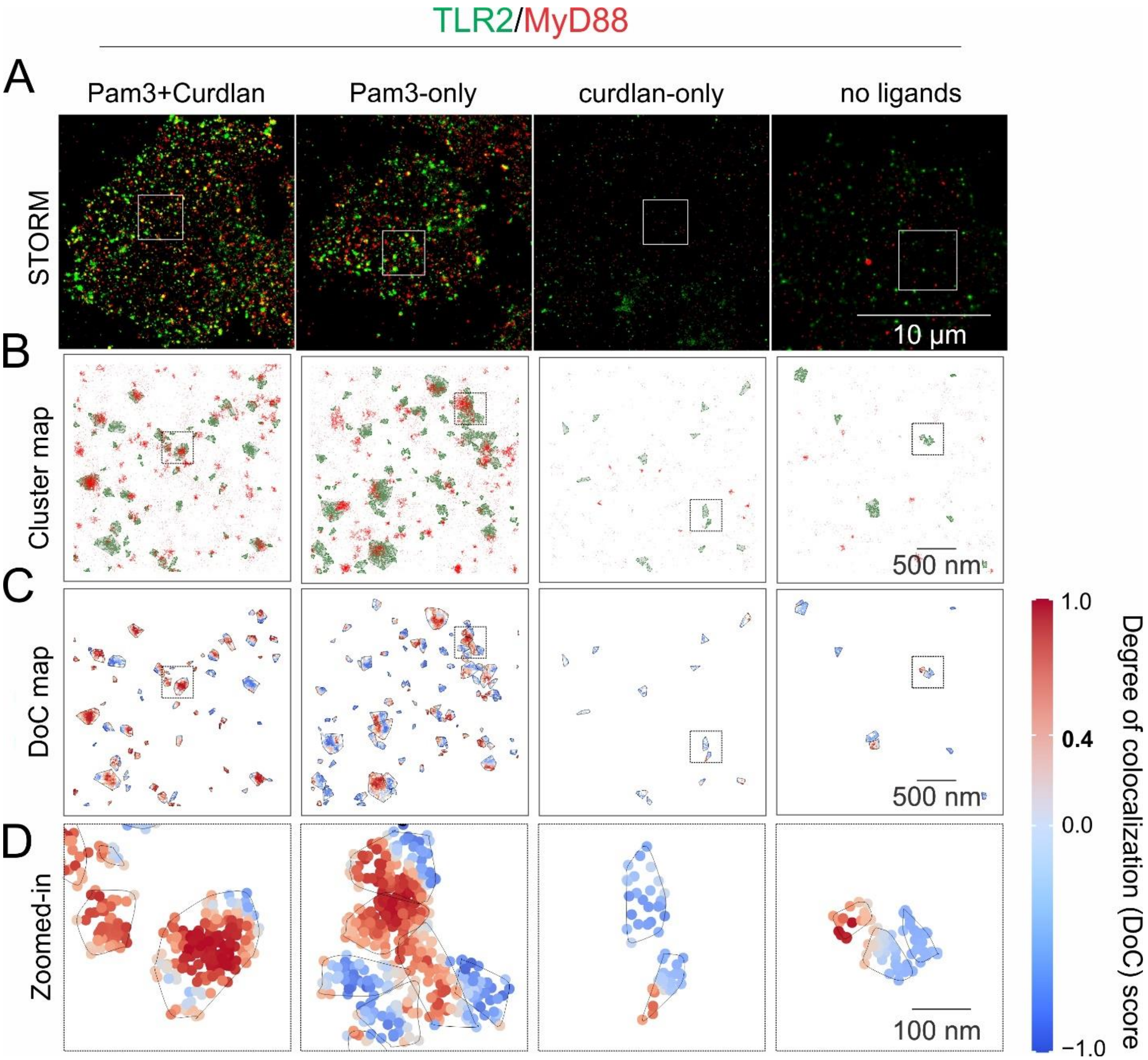


**Fig. S17: Quantification of the activation of TLR2 nanoclusters. (A)** Representative two-color dSTORM images of immunofluorescently labeled TLR2 and MyD88 in RAW264.7 macrophage cells seeded on various substrates for 10 min. **(B)** Cluster maps of TLR2 (green dots) and MyD88 (red dots) from the zoomed-in areas outlined in **(A)**. **(C, D)** Degree of colocalization (DoC) analysis of TLR2 that overlapped with MyD88. Single molecule localizations of TLR2 with DoC ≥ 0.4 are considered as activated. Receptor nanocluster contours are shown in black lines.

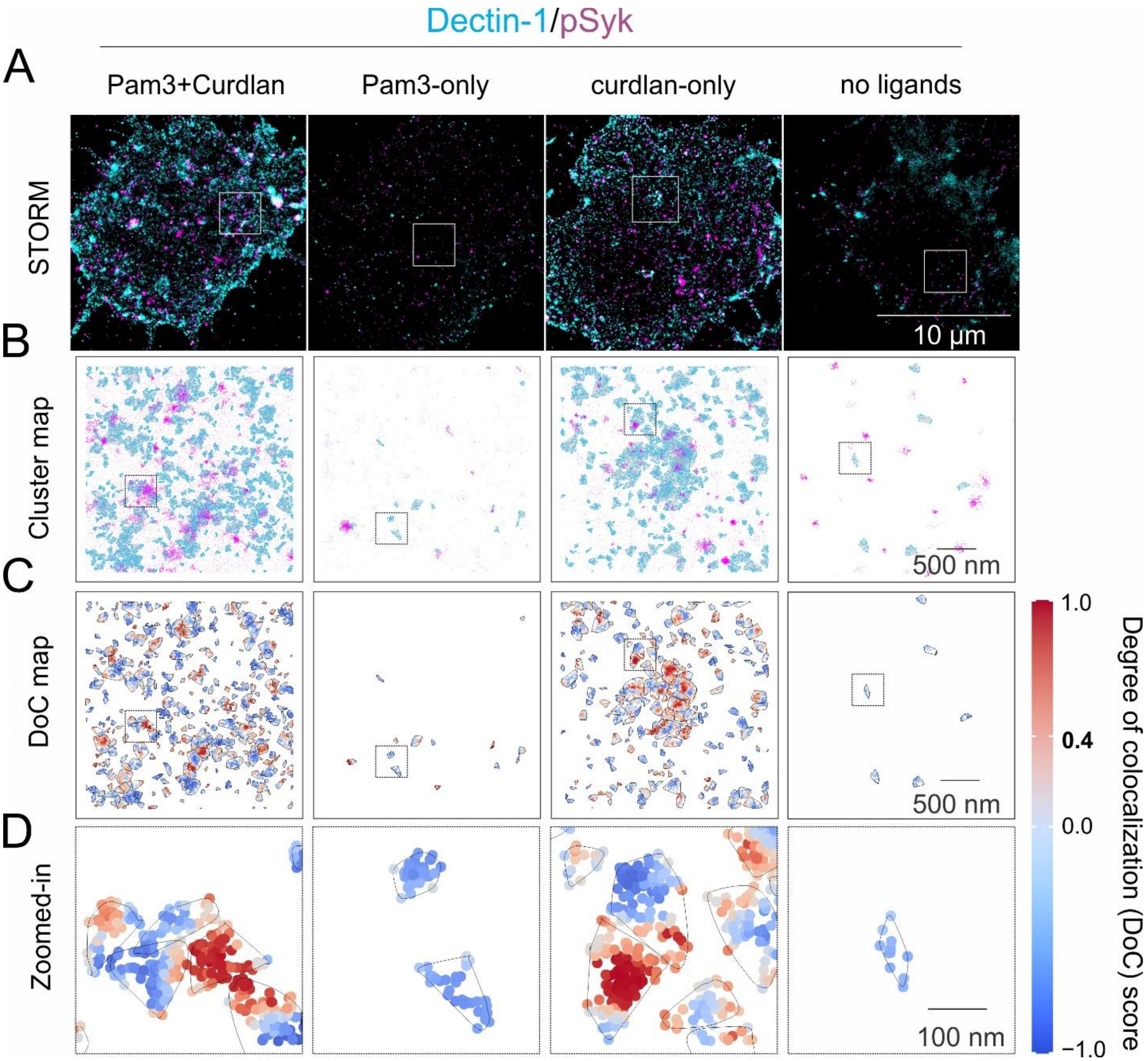


**Fig. S18: Quantification of the activation of Dectin-1 nanoclusters. (A)** Representative two-color dSTORM images of immunofluorescently labeled Dectin-1 and pSyk in RAW264.7 cells seeded on various substrates for 10 min. **(B)** Cluster maps of Dectin-1 (cyan) and pSyk (magenta) from zoomed-in areas outlined in **(A)**. **(C, D)** Degree of colocalization (DoC) analysis of Dectin-1 that overlapped with pSyk. Single molecule localizations of Dectin-1 with DoC ≥ 0.4 are considered as activated. Receptor nanocluster contours are shown in black lines.
